# Spectral fiber-photometry derives hemoglobin concentration changes for accurate measurement of fluorescent sensor activity

**DOI:** 10.1101/2021.08.23.457372

**Authors:** Wei-Ting Zhang, Tzu-Hao Harry Chao, Yue Yang, Tzu-Wen Wang, Sung-Ho Lee, Esteban A. Oyarzabal, Jingheng Zhou, Randy Nonneman, Nicolas C. Pegard, Hongtu Zhu, Guohong Cui, Yen-Yu Ian Shih

## Abstract

Fiber-photometry is an emerging technique for recording fluorescent sensor activity in the brain. However, significant hemoglobin-absorption artifacts in fiber-photometry data may be misinterpreted as sensor activity changes. Because hemoglobin exists in nearly every location in the brain and its concentration varies over time, such artifacts could impede the accuracy of many photometry recording results. Here we present a novel use of spectral photometry technique and propose computational methods to quantify photon absorption effects by using activity-independent fluorescence signals, which can be used to derive oxy- and deoxy-hemoglobin concentration changes. Following time-locked neuronal activation *in vivo*, we observed that a 20% increase in CBV contributes to about a 4% decrease in green fluorescence signal and a 2% decrease in red fluorescence signal. While these hemoglobin concentration changes are often temporally delayed than the fast-responding fluorescence spikes, we found that erroneous interpretation may occur when examining pharmacology-induced sustained activity changes, and in some cases, hemoglobin-absorption could flip the GCaMP signal polarity. We provided hemoglobin-based correction methods to restore fluorescence signals across spectra and compare our results against the commonly used regression approach. We also demonstrated the utility of spectral fiber-photometry for delineating brain regional differences in hemodynamic response functions.

**Highlights:**

- Hemoglobin-absorption compromises fiber-photometry recording *in vivo*
- Spectral photometry allows quantification of hemoglobin concentration changes for correction
- The proposed platform allows measuring regional differences in neurovascular transfer function

## Introduction

Fiber-photometry relies on an implanted optical fiber to excite and detect fluorescence signals, and is best known for its ability to measure ensemble neuronal or neurochemical activity in small volumes of brain tissue in animal models. The simplicity and cost-effectiveness of the optical system make the technique highly accessible (Meng et al., 2018), and lightweight, miniature fiber optics make photometry compatible with a wide variety of experimental settings, such as freely moving behavioral assessment (Kim et al., 2016; Meng et al., 2018) or head-fixed brain-wide imaging (Chen et al., 2019; Pais-Roldán et al., 2020) (**Figure 1A**). Several genetically encoded fluorescent sensors of cellular activity and neurotransmitters have recently become widely available (Chen et al., 2013; Dana et al., 2016; Marvin et al., 2019, 2013; Patriarchi et al., 2018; Sun et al., 2018; Tian et al., 2009; H. Wang et al., 2018), expanding the range of applications for fiber-photometry. The tunable spectral specificity has further enabled new parallel sensing capabilities with wavelength multiplexing. Given its rapidly increasing usage in neuroscience (Cui et al., 2013; Luchsinger et al., 2021; Pisano et al., 2019; Sych et al., 2019), understanding the inherent confounding factors in fiber-photometry data is of paramount significance.

**Figure 1.**
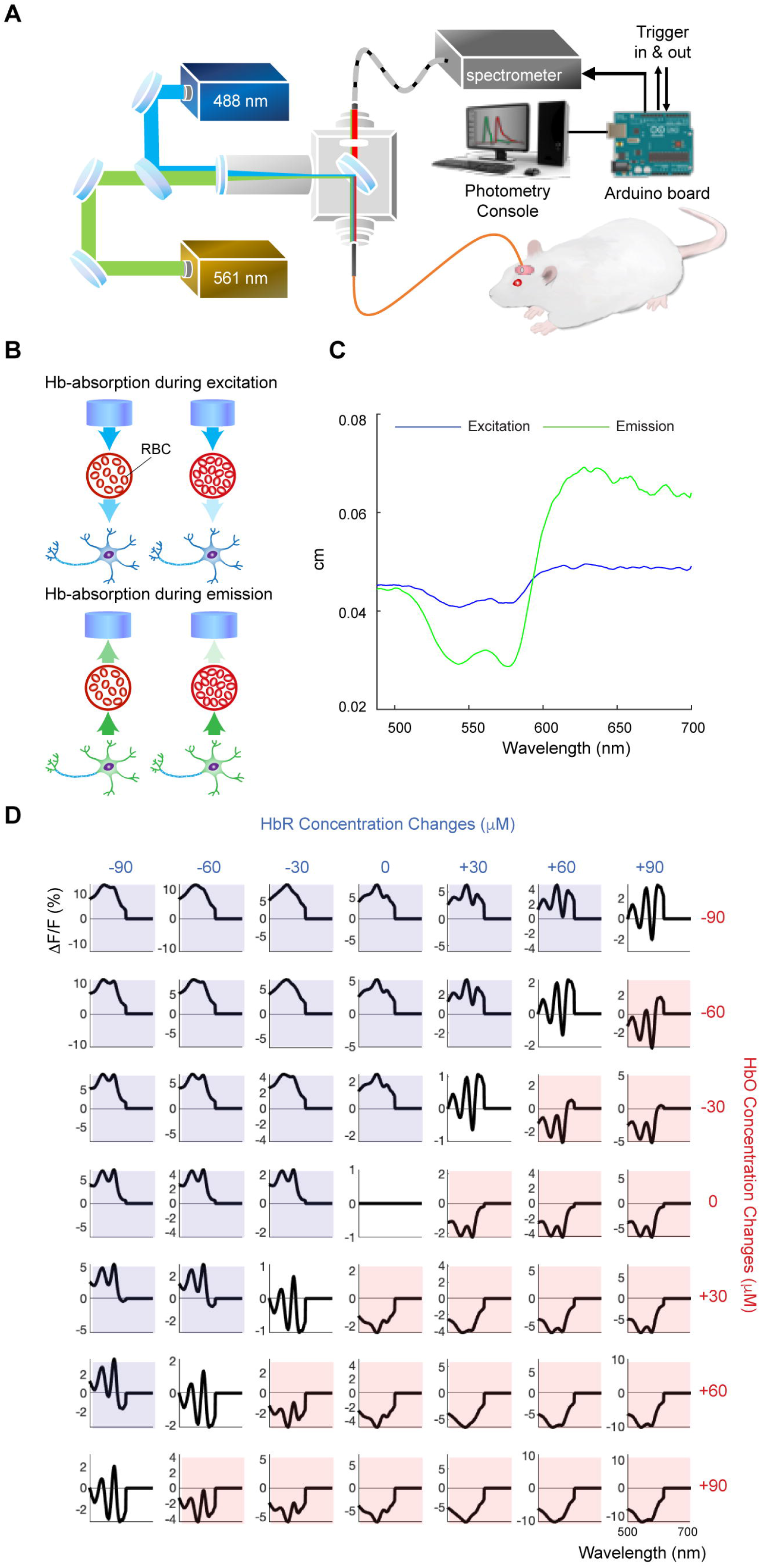
Spectrally resolved fiber-photometry system and Hb-absorption effects. (**A**) Schematic illustration of a spectrally resolved fiber-photometry system used for recording fluorescence signals. (**B**) Hb in the red blood cells absorb light in two phases: the excitation light from the optical fiber (left) and the emission light to be recorded by the optical fiber (right). Color density of arrowheads delineates light intensity. Increase in Hb is expected to decrease the number of photons in both phases. (**C**) Simulated average excitation and emission photon traveling pathlengths across 488 – 700 nm emission wavelengths from our database of paired excitation/emission light paths. (**D**) Impact of HBO and HBR concentration on ΔF/F. Simulated effects of HbO and HbR concentration changes on fluorescence signal measurement. Δ*C_Hb0_* and Δ*C_HbR_* ranging from −90 to +90 μM were used to simulate fluorescence signal changes (ΔF/F) between 500 to 700 nm using the molar extinction coefficients of HbO and HbR in **Figure S1** and simulated light paths in **Figure 1C**. Graphs shaded in blue indicate an overall decrease in total Hb (HbT); whereas graphs shaded in red indicate an overall increase in HbT. See additional details in Figure S1–S4, Supplementary Video 1, and Table S1.

One of the most notable confounders affecting fluorescence measurements is light absorption by hemoglobin (Hb) within blood vessels (Ma et al., 2016; Prahl, 1999). The Hb-absorption artifact has been studied and addressed extensively in wide-field imaging with reflective or backscattered lights (Ma et al., 2016; Valley et al., 2020) and recently in two-photon lifetime microscopy (2PLM) measuring the partial pressure of O_2_ (Mächler et al., 2021). The Hb-absorption artifact should be present in most photometry data because the intervascular distance of capillaries is known to be < 60 *μm* in nearly every location in the brain (Smith et al., 2019) and commonly used photometry targeted volumes is 10^5^–10^6^ *μm*^3^ (Pisanello et al., 2019). This absorption occurs in two phases: along the excitation path, by absorbing photons that travel from the fiber tip to the targeted fluorescent proteins; and along the emission path, by absorbing photons emitted by the fluorescent proteins as they travel back towards the optical fiber (**Figure 1B**). Both absorption effects add up and result in a substantial photon count reduction. Importantly, Hb-absorption is non-linear, wavelength-dependent, and varies as a function of dynamic oxy- and deoxy-hemoglobin (HbO and HbR) concentration changes *in vivo* (Δ*C_HbO_* and Δ*C_HbR_*) (Meng and Alayash, 2017; Prahl, 1999) (**Figure S1**, derived from(Prahl, 1999)), causing erroneous photometry reading. Further, because the power spectra of hemodynamic signals overlap significantly with most commonly used fluorescent sensors (**Figure S2**), hemoglobin related artifacts cannot be removed from fiber-photometry data by notch filtering or regressing either cerebral blood flow or cerebral blood volume (CBV) signals. To the best of our knowledge, the influence of Hb-absorption on fiber-photometry-derived fluorescent sensor activity has never been studied. The ability to quantify Hb concentration fluctuations in photometry data would not only enable more accurate fluorescent sensor activity measurements, but would also provide concurrent hemodynamic metrics, which is the foundation of several brain-mapping techniques including functional magnetic resonance imaging (fMRI) (Schulz et al., 2012), intrinsic optics (Hillman, 2007), and functional ultrasound (Rungta et al., 2017).

In this study, we propose a novel use of spectrally resolved fiber-photometry platform and computational methods that can independently derive HbO and HbR concentration changes from activity-independent fluorescent reporter signals. We present the use of this technique to remove the Hb-absorption artifacts from calcium activity recording and demonstrate the added benefit to compute transfer function between neuronal and vascular activity which is crucial for interpreting hemodynamic-based neuroimaging data such as fMRI, functional ultrasound, and intrinsic optics.

## Results

### *Theoretical Hb-absorption* effect on photometry recordings

We solved the following equation using the generalized method of moments (GMM) (Hansen, 1982) with spectrometer outputs, which provides quantifiable photon counts across spectra:

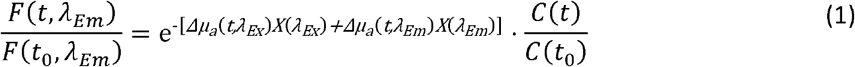

where *Δμ_a_*(*t,λ_Ex_*) and *Δμ_a_*(*t,λ_Em_*) represent the absorption coefficient changes at excitation wavelength *λ_Ex_* and emission wavelength *λ_Em_*, respectively, at time point *t* as compared to a reference time point *t*_0_. Both *Δμ_a_*(*t,λ_Ex_*) and *Δμ_a_*(*t,λ_Em_*) are affected by Δ*C_HbO_* and Δ*C_HbR_* (see Method Equation 16). *X*(*λ_Ex_*) and *X*(*λ_Em_*) represent the photon traveling pathlengths (see Methods for details) at the excitation and emission wavelengths, respectively. To solve equation (1), we first performed a Monte Carlo simulation using the pipeline shown in **Figure S3** to obtain *X*(*λ_Ex_*) and *X*(*λ_Em_*). We determined the average photon traveling pathlengths for emission wavelengths from 488 nm to 700 nm in the simulated brain tissue from a total of 29 simulations (2-2.5 × 10^4^ photons each) in **Figure 1C**, with detailed numbers tabulated in **Table S1**. These results are expected to be generalizable and can be used across labs. As an example, we illustrated spatial probability and luminance-distance profiles for the commonly used 488 nm excitation, 515 nm emission and 580 nm emission in **Figure S4A-C** (see also **Supplementary Video** 1). These results allowed calculating the simulated effects of Hb-absorption on fluorescence signal measurement, for which we observed significant nonlinear, wavelength-dependent effects (**Figure 1D**).

### Ex vivo quantification of Hb-absorption effects on photometry recordings

We first performed *ex vivo* measurements and showed that adding 1 mL of fresh, venous mouse blood into a 10 mL mixture of Alexa 488 and 568 fluorescent dye solution significantly reduced both fluorescence signals measured by fiber-photometry. Compared to adding the blood, adding 1 mL vehicle (0.9% saline) induced significantly less reduction in both Alexa 488 and 568 signals (**Figure S5A, C**). Subsequently, we saturated the mixture with 100% oxygen to increase the HbO-HbR ratio and observed marked increases in the Alexa 568 signals but decreases in Alexa 488 signals. We did not observe significant signal changes when infusing the oxygen into the solution added with 1 mL saline (**Figure S5B, D**).

### In vivo quantification of Hb-absorption effects on photometry recordings

We performed an *in vivo* experiment to measure the Hb-absorption of activity-independent fluorescence signals in fiber-photometry. We selected a commonly used fluorophore, enhanced yellow fluorescent protein (EYFP), whose fluorescence signal is activity-independent and has an emission spectrum peaking at 526 nm. We virally expressed EYFP using adeno-associated virus (AAV) in the primary somatosensory cortex (S1) of the forelimb region (S1FL) under the calmodulin kinase IIα (CaMKIIα) promoter and then implanted an optical fiber into this target area (**Figure 2A, B**). On the day of photometry recording, we intravenously administered Rhodamine B, a long-circulating red fluorescent dye, to enable steady-state CBV measurements (Unekawa et al., 2015) (**Figure 2C**) and cross validate our proposed method in calculating total Hb concentration changes. Of note, the resulting EYFP and Rhodamine B spectra overlapped significantly with the Hb-absorption spectra (see **Figure S2**), suggesting a significant impact of Hb-absorption on the data from these fluorescence signals. Upon electrical stimulation of the contralateral forepaw (9 Hz, 3 mA, 0.5 ms pulse width for 10 s), we observed robust positive Rhodamine-derived CBV (rhoCBV) changes as expected (**Figure 2D**). Intriguingly, the EYFP signal concurrently recorded from the same fiber showed robust negative changes (**Figure 2E**), which indicated significant Hb-absorption of the EYFP signal *in vivo*. Although the pH-dependent effect on the EYFP signal cannot be completely excluded, a previous study showed that in pH 6.5-9, where typical pH homeostasis of a hippocampal neuron (^~^7.03-7.46) is located (Ruffin et al., 2014), EYFP excited by 470 nm was not significantly affected by the pH changes (Nakabayashi et al., 2012). Using well-documented molar extinction coefficients (**Figure S1**), photon-traveling pathlengths derived from a Monte Carlo simulation (**Figure 1C**), and spectral fiber-photometry datapoints across 502 – 544 nm wavelengths over time (see equation (17–21) in **Methods**), we derived Δ*C_HbO_*(*t*) and Δ*C_HbR_*(*t*) using GMM. Notably, this is feasible and robust because we have access to a range of spectral datapoints for every single time point and therefore could use numerous empirical data to solve two unknowns (Δ*C_HbO_* and Δ*C_HbR_*, see also **Figure S6** for the effect of input spectral datapoints). The derived concentration changes (**Figure 2F, G**) were within physiological ranges and comparable to those reported in the literature (Ma et al., 2016), and the summation of the two (i.e., total Hb concentration changes, termed as Δ*C_HbT_*(*t*)) informs CBV changes (Ma et al., 2016) (**Figure 2J**). The calculated Δ*C_HbT_*(*t*) highly resembled the rhoCBV changes measured from a distinct and less-affected red spectrum (**Figure S7**). With Δ*C_HbO_*(*t*) and Δ*C_HbR_*(*t*) obtained, we then calculated *Δμ_a_*(*t,λ_Ex_*) and *Δμ_a_*(*t,λ_Em_*), and input these parameters into equation (1) to correct the rhoCBV changes (**Figure 2H**) and EYFP signals (**Figure 2I**). Indeed, **Figure 2I** shows a flattened EYFP trace following the proposed correction, demonstrating the validity of our approach. Although the influence of Hb-absorption on EYFP signal varied by trials, our proposed method accurately corrected EYFP signals in all trials (**Figure 2K**). Notably, despite the smaller influence on the rhoCBV signal measured from the red spectrum, the absorption effect remained statistically significant (**Figure 2L**). In addition to the fluorescence signal changes induced by forepaw sensory stimulations, we also replicated these findings using a 5% CO_2_ hypercapnic challenge paradigm and successfully restored the contaminated EYFP signal changes (**Figure S8**).

**Figure 2.**
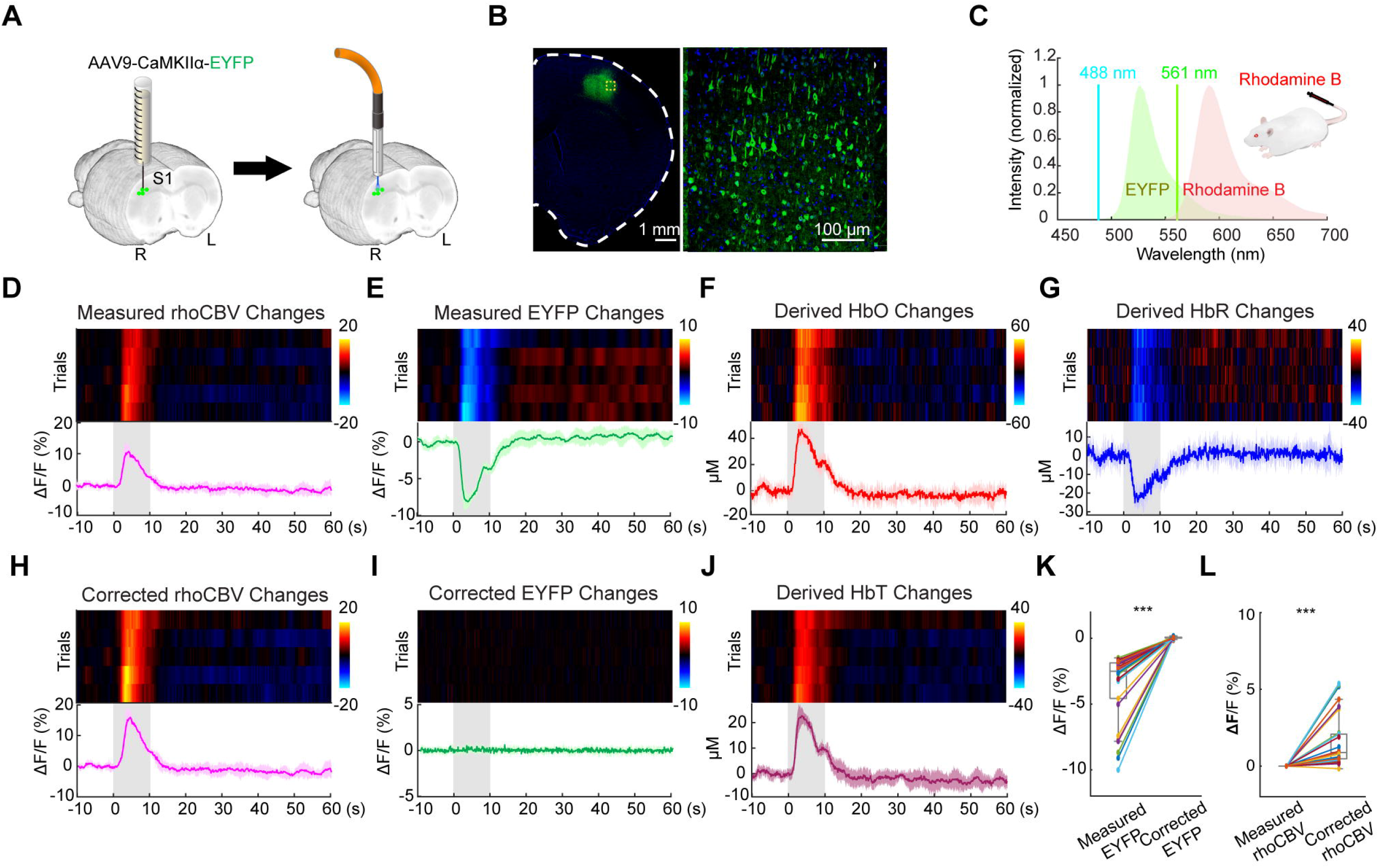
In vivo quantification of HbO and HbR changes using EYFP emission in fiber-photometry and correction of EYFP signals for Hb-absorption. (**A**) Preparation for fiber-photometry measurement of EYFP signals in the S1 of the forelimb region. (**B**) Confocal image showing EYFP expression. (**C**) Simultaneous dual-spectral photometry recording of EYFP signals and hemodynamic responses (CBV) was achieved by intravenously injecting a long-circulating red fluorescent dye, Rhodamine B. (**D-J**) Peri-stimulus heatmaps showing repeated trials and average response time-courses of the changes in measured rhoCBV, measured EYFP, derived HbO, derived HbR, corrected rhoCBV, corrected EYFP, and derived HbT, respectively. The gray-shaded segment indicates forepaw-stimulation period. Note that Hb-absorption induce a pseudo-negative changes in measured EYFP signals during stimulation. (**K,L**) Correction of the contaminated EYFP and rhoCBV signals, respectively, across 5 subjects and 30 trials (\*\*\**p* < 0.001, paired *t*-test). Measured and corrected EYFP signals are compared. Measured rhoCBV was normalized to 0 to show the difference before and after the correction. All color bars use the same unit as the time-course Y axis. Error bars represent standard deviation. See also Figure S6–S8.

### Impact of Hb-absorption on acute GCaMP signal measurement using fiber-photometry

We virally expressed the GCaMP6f (hereafter as GCaMP), one of the most widely used genetically encoded calcium indicators (Chen et al., 2013), using AAV under the CaMKIIα promoter in the S1FL. We also expressed tandem dimer tomato (tdTomato), an activity-independent red fluorescent protein, under the CAG promoter (**Figure 3A, B**) and then implanted an optical fiber into this target area. Our rationale for choosing tdTomato was similar to that for EYFP in **Figure 2**, as it does not encode activity changes and thus its photon counts can be used to derive Δ*C_HbO_* and Δ*C_HbR_* for GCaMP signal correction. Using different promoters for GCaMP and tdTomato doesn’t affect the application of our method in that tdTomato serves as “background light” which is irrelevant to the neuronal changes but is generally affected by the vascular changes no matter which promoter was chosen. Unlike in the experiment described in **Figure 2**, Rhodamine B could no longer be used for CBV cross-validation because the red spectrum was occupied by tdTomato (**Figure 3C**). Instead, we performed this GCaMP-tdTomato experiment inside a magnetic resonance imaging (MRI) scanner where CBV changes could be measured by intravenously administering an iron oxide contrast agent, Feraheme^23^ (**Figure 3D**). This allowed CBV to be validated using an established and completely independent modality. After a short, 1 s electrical stimulation of the contralateral forepaw (9 Hz, 3 mA, 0.5 ms pulse width), we observed a rapid increase in GCaMP signal followed by an undershoot (**Figure 3E, Q**). Such undershoot is absent in cultured neurons where Hb is not present and rarely observed in two-photon imaging data where neurons and blood vessels are mostly segregated in space (Chen et al., 2013; Mittmann et al., 2011). The tdTomato signal decreased significantly upon forepaw stimulation and showed a much longer delay in its kinetics, but has its peak temporally matched to that of the GCaMP undershoot (**Figure 3F**). Using the methods similar to that described in **Figure 2**, but with spectral photometry datapoints across 575-699 nm wavelengths over time, we derived Δ*C_HbO_*(*t*) and Δ*C_HbR_*(*t*) (**Figure 3G, H**). Interestingly, the derived ΔC_HbR_ data exhibited a rapid initial increase (**Figure 3H**) before a more prolonged reduction – a phenomenon well-documented in the optical literature(Devor et al., 2003) and termed “initial dip” in blood-oxygenation-level-dependent fMRI(Hu and Yacoub, 2012). This initial increase in HbR was not detectable in the raw tdTomato data, indicating that our method is capable of deriving parameters with temporal kinetics distinct from the detected raw fluorescence activity. **Figure 3J** shows the corrected GCaMP signals, where the after-stimulus undershoot no longer exists. Accordingly, **Figure 3K** shows a flattened tdTomato trace following the correction. The derived Δ*C_HbT_*(*t*) (**Figure 3I**) highly resembled the CBV changes measured by fMRI (**Figure 3L**) and these two time-courses correlated significantly (**Figure 3M**). The derived Δ*C_HbT_* showed higher sensitivity than MRI-CBV (**Figure 3N**). We observed no significant time-lag between the Δ*C_HbT_* and CBV-fMRI time-courses (**Figure 3O, P**).

**Figure 3.**
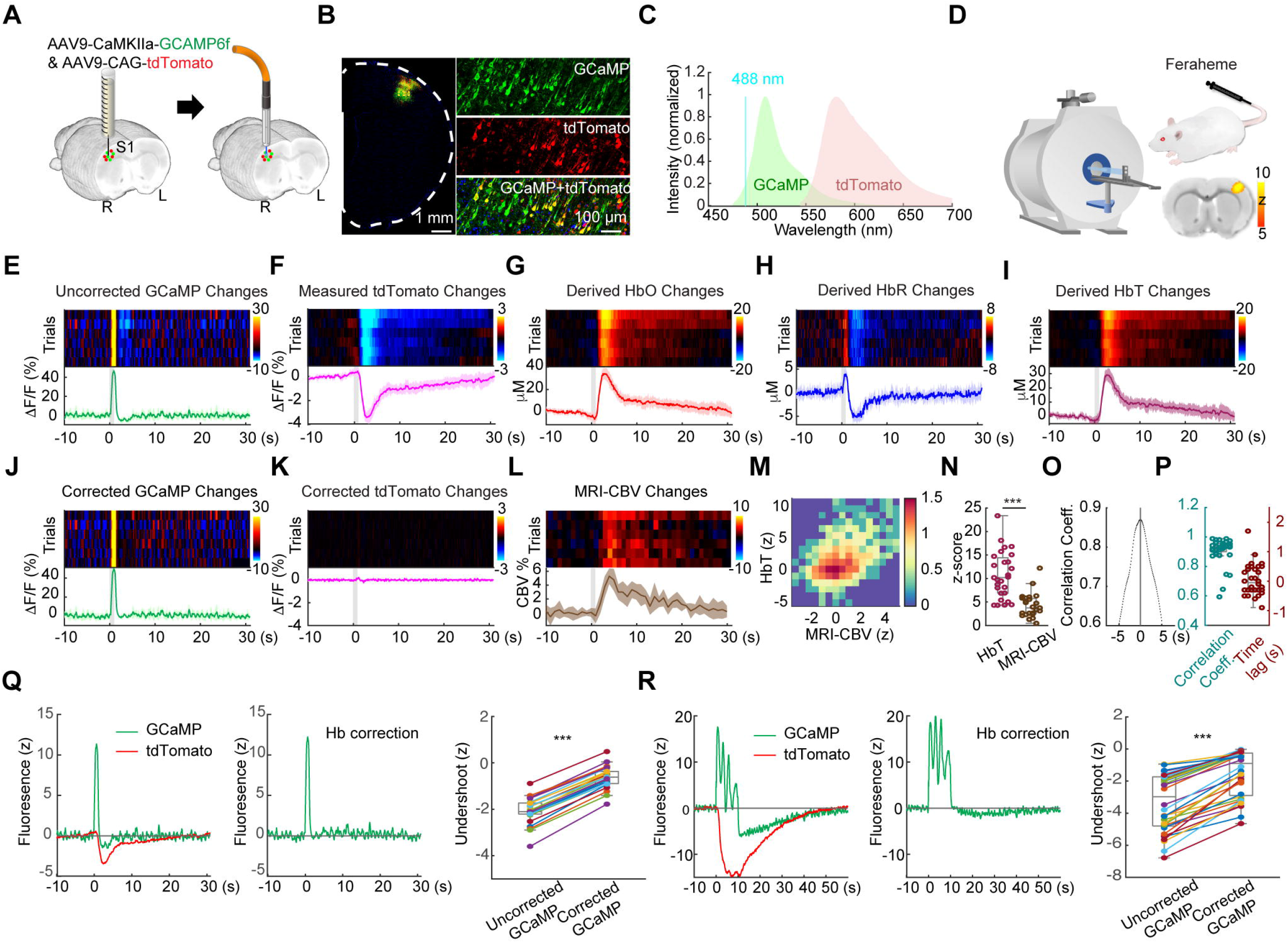
In vivo quantification of HbO and HbR changes using tdTomato emission in fiber-photometry and correction of GCaMP signals for Hb-absorption. (**A**) Preparation for fiber-photometry measurement of GCaMP and tdTomato signals in the S1 of the forelimb region. (**B**) Confocal images showing GCaMP and tdTomato expression. (**C**) GCaMP and tdTomato emission spectra from 488 nm excitation. (**D**) Concurrent CBV measurement was achieved by MRI and intravenous injection of the long-circulating iron oxide nanoparticles, Feraheme. Insert shows a forepaw-induced group-level fMRI activation map (corrected p < 0.05, n = 5). (**E-L**) Peri-stimulus heatmaps showing repeated trials and average response time-courses of the changes in measured GCaMP, measured tdTomato, derived HbO, derived HbR, derived HbT, corrected GCaMP, corrected tdTomato, and measured MRI-CBV, respectively. The gray-shaded segment indicates the forepaw-stimulation period. (**M**) Two-dimensional histogram summarizing HbT and MRI-CBV changes across 5 subjects and 27 trials. The color bar indicates counts on the log scale. (**N**) Photometry derived HbT exhibited higher sensitivity than MRI-CBV (\*\*\**p* < 0.001, paired *t*-test). (O) Trial-level cross-correlation between the Z scores of the derived HbT and the Z scores of the MRI-CBV over the peri-stimulus time period of 10 s, and (**P**) scatter plots showing cross-correlation coefficients (left) and time lag (right) of all trials. (**Q, R**) Correction of pseudo-undershoot artifacts in GCaMP time-courses of (**Q**) short (1 s), and (**R**) long (10 s) stimulations, respectively ((\*\*\**p* < 0.001, paired *t*-test). See also Figure S9–S10.

**Figure 3Q and 3R** demonstrate the corrections of pseudo-undershoot artifacts in GCaMP timecourses of short (1 s) and long (10 s) stimulations, respectively. These artifacts following short bursts of activation were also apparent when optogenetic stimulation was applied in the brain (**Figure S9**). In addition, we also observed a significant correction effect on GCaMP response amplitudes (**Figure S10**). Without proper correction of Hb-absorption artifacts, the accuracy of quantified fluorescence signals is compromised, leading to misinterpretation of fiber-photometry data. This problem could become exacerbated in the absence of time-locked stimulations, as the desired activity could be superimposed on physiological hemodynamic signal changes.

### Impact of Hb-absorption on sustained GCaMP signal measurement using fiber-photometry

To further demonstrate the impact of Hb-absorption on GCaMP signal over an extended period of time, we leverage an acute ethanol challenge protocol (Broadwater et al., 2018; Vaagenes et al., 2015) which has been shown to decrease neuronal activity (Lotfullina and Khazipov, 2018) as well as cerebral perfusion (Broadwater et al., 2018; Kelly et al., 1997; Sano et al., 1993). We virally expressed GCaMP using AAV under the CaMKIIα promoter and tdTomato under the CAG promoter in the prelimbic cortex (PrL) (**Figure 4A**) and then implanted an optical fiber into this target area. On the day of the experiment, ethanol (2 g/kg, 20% v/v, i.p.) was given 5 min after the onset of fiber-photometry recording. After ethanol administration, we observed robust increases in the raw, measured tdTomato (**Figure 4B**) and GCaMP (**Figure 4C**) signals. These changes were ethanol specific, as we did not observe any changes in a separate session where the same group of subjects received vehicle injection (an equal volume of 0.9% saline, i.p.) (**Figure 4B, C**). Next, we derived Δ*C_HbO_(t)*, Δ*C_HbR_(t)* and Δ*C_HbT_(t)* (**Figure 4D**) from the ethanol data using the correction methods identical to that described in **Figure 3** and found robust decreases in Δ*C_HbO_(t)* and Δ*C_HbT_ (t)*, in agreement with studies showing reduced Hb affinity for oxygen (Van De Borne et al., 1997) and vasoconstriction (Kudo et al., 2015) following acute alcohol administration. Following our proposed correction, tdTomato trace was flattened as expected (**Figure 4B, E**). Importantly, the trend of the GCaMP signal was flipped from the unexpected positive changes to the expected negative changes (Lotfullina and Khazipov, 2018) (**Figure 4C, F**). In another two proof-of-concept experiments, we also noted the influence of Hb-absorption on anesthetics- and pharmacology-induced fluorescence signal changes (**Figures S11 and S12**). These data highlight the importance to correct for Hb-absorption in fiber-photometry studies, as misinterpretation may otherwise occur.

**Figure 4.**
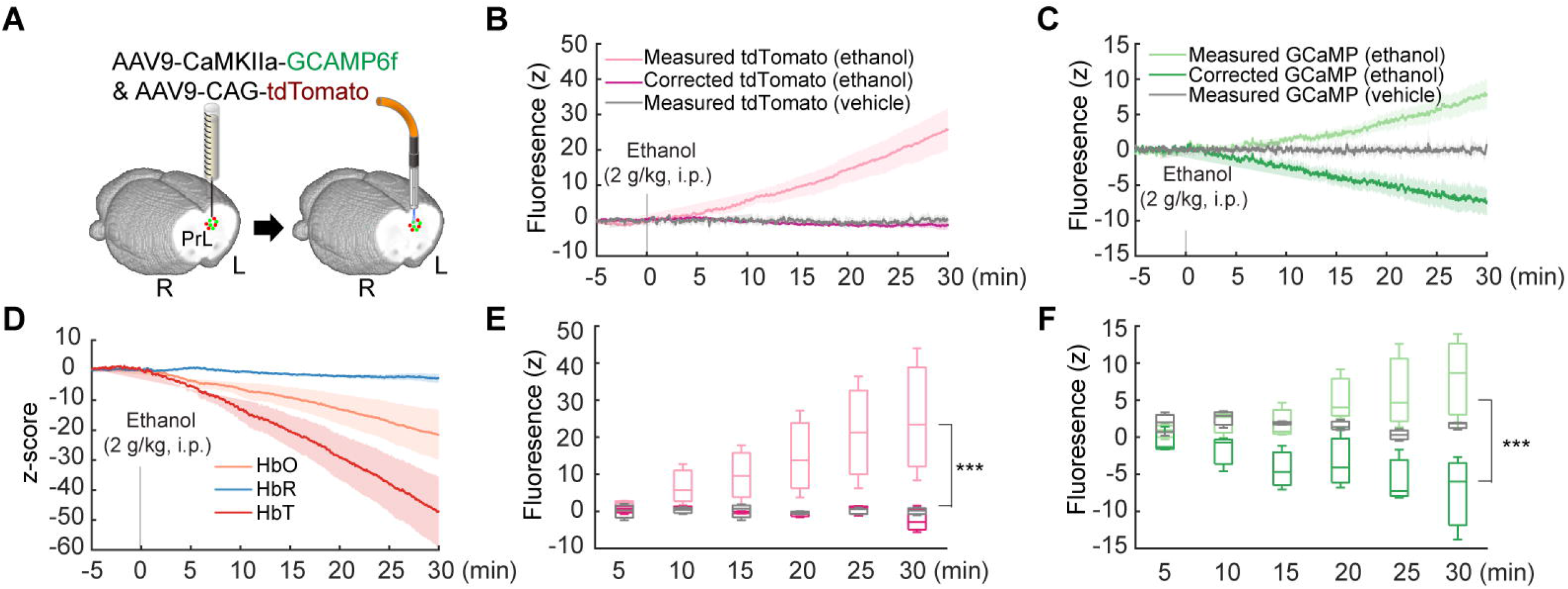
Hb-absorption effects on sustained GCaMP signal measurement using fiber-photometry. **(A)** Preparation for fiber-photometry measurement of GCaMP and tdTomato signals in the PrL region. **(B)** Measured and corrected tdTomato time-courses before and after ethanol administration. The same volume of saline was administered in a separate session to generate within-subject vehicle control data. **(C)** Measured and corrected tdTomato signals were sampled and displayed at 5 min. A significant effect was found between the three groups and *post-hoc* test showed a significant difference between the measured and corrected signals. \*\*\**p* < 0.001, two-way ANOVA (n=3). **(D)** Derived HbO, HbR, and HbT time-courses before and after the ethanol administration. Higher HbT z-scores were attributed to the lower baseline variation following the addition of HbO and HbR. **(E)** Measured and corrected GCaMP time-courses before and after the ethanol administration. The same volume of saline was administered in a separate session to generate within-subject vehicle control data. Data were collected concurrently with those shown in (B). **(F)** Measured and corrected GCaMP signals were sampled and displayed at 5 min intervals. A significant effect was found between the three groups and *post-hoc test* showed a significant difference between the measured and corrected signals. \*\*\**p* < 0.001, two-way ANOVA (n=3).

### Correction of Hb-absorption artifacts in other fiber-photometry settings

To expand the utility of our method, we modeled and compared the Hb-absorption correction techniques among the following three settings: **(1)** Using a spectrometer with a sequence that interleaves between 405 and 488 nm excitation. GCaMP isosbestic point is around 405 nm (Tian et al., 2009; Dana et al., 2019), where excitation light delivered at this range would evoke GCaMP emission photons irrelevant to the changes in calcium concentration. The GCaMP emission signals from the 405 nm excitation is abbreviated as the GCaMP_405_, in contrast to the GCaMP_488_ excited by the 488 nm laser. The fact that apparent Hb-absorption exists at GCaMP_405_ emission spectrum (**Figure S1**) makes this an appealing alternative approach when another activity-independent fluorescent protein is not expressed. In essence, this provides the reference fluorescence signals similar to that of EYFP in **Figure 2** or tdTomato in **Figure 3**, and allows Δ*C_HbO_ (t)* and Δ*C_HbR_ (t)* to be calculated for Hb-absorption correction; **(2)** using GCaMP_405_ data from a spectrometer but only selected three spectral datapoints to mimic the settings using band-pass filters with a limited number of photodetectors (i.e., without using the full GCaMP_405_ spectrum for correction); and (3) regressing out the activity-independent GCaMP_405_ fluorescence signal intensity as seen in (Kim et al., 2016). To provide data for these evaluations, we virally expressed GCaMP6f using AAV under the CaMKIIα promoter and tdTomato under the CAG promoter in the primary motor cortex (M1) of the left hemisphere, and implanted a microelectrode into the M1 of the right hemisphere, allowing electrical stimulation to be delivered (60 Hz, 300 μA, 0.5 ms pulse width for 2 s) and thus time-locked GCaMP signal changes can be induced in the left M1 (**Figure 5A**). The interleaved excitation and acquisition sequences are shown in **Figure 5B**. The raw GCaMP_405_ and GCaMP_488_ responses to stimulation are shown in **Figure 5C**. Similar to that in **Figure 3Q & R**, we observed a rapid increase in the GCaMP_488_ signal, followed by a pseudo-undershoot (Chen et al., 2013; Mittmann et al., 2011). We then compared the aforementioned three settings using this single set of *in vivo* data, thus avoiding a subject and trial-by-trial variability confounds. Respectively to the aforementioned three settings, we present: **(1)** corrected GCaMP_488_ using Δ*C_HbO_ (t)* and Δ*C_HbR_ (t)* derived from the full GCaMP_405_ spectrum in **Figure 5D, (2**) corrected GCaMP_488_ using Δ*C_HbO_ (t)* and Δ*C_HbR_ (t)* derived from three spectral datapoints from the GCaMP_405_ spectrum in **Figure 5E**, and **(3)** corrected GCaMP_488_ by regressing out the GCaMP_405_ signal intensity in **Figure 5F**. Comparisons of the post-stimulus pseudo-undershoot artifacts suggested successful restoration of the GCaMP_488_ signals (i.e., signals rapidly return to zero) via both Hb-absorption-based correction methods, but not via the regression-based method (**Figure 5G**). As tdTomato signals are available from 488 nm excitation (**Figure 5B**), we carried out a similar set of analyses by using tdTomato instead of GCaMP_405_ emission signals (**Figures 5H-K**) and observed successful restoration of the GCaMP_488_ signals via both Hb-absorption-based correction methods, but sustained elevated signals by the regression-based method (**Figures 5H-K**). No significant difference was observed between the full spectrum vs. three-point correction methods (**Figures 5G & L**). Nevertheless, as expected, the HbT computed from either GCaMP_405_ or tdTomato spectrum exhibited lower CNR when using only three spectral datapoints compared to the full spectrum (**Figure 5M**).

**Figure 5.**
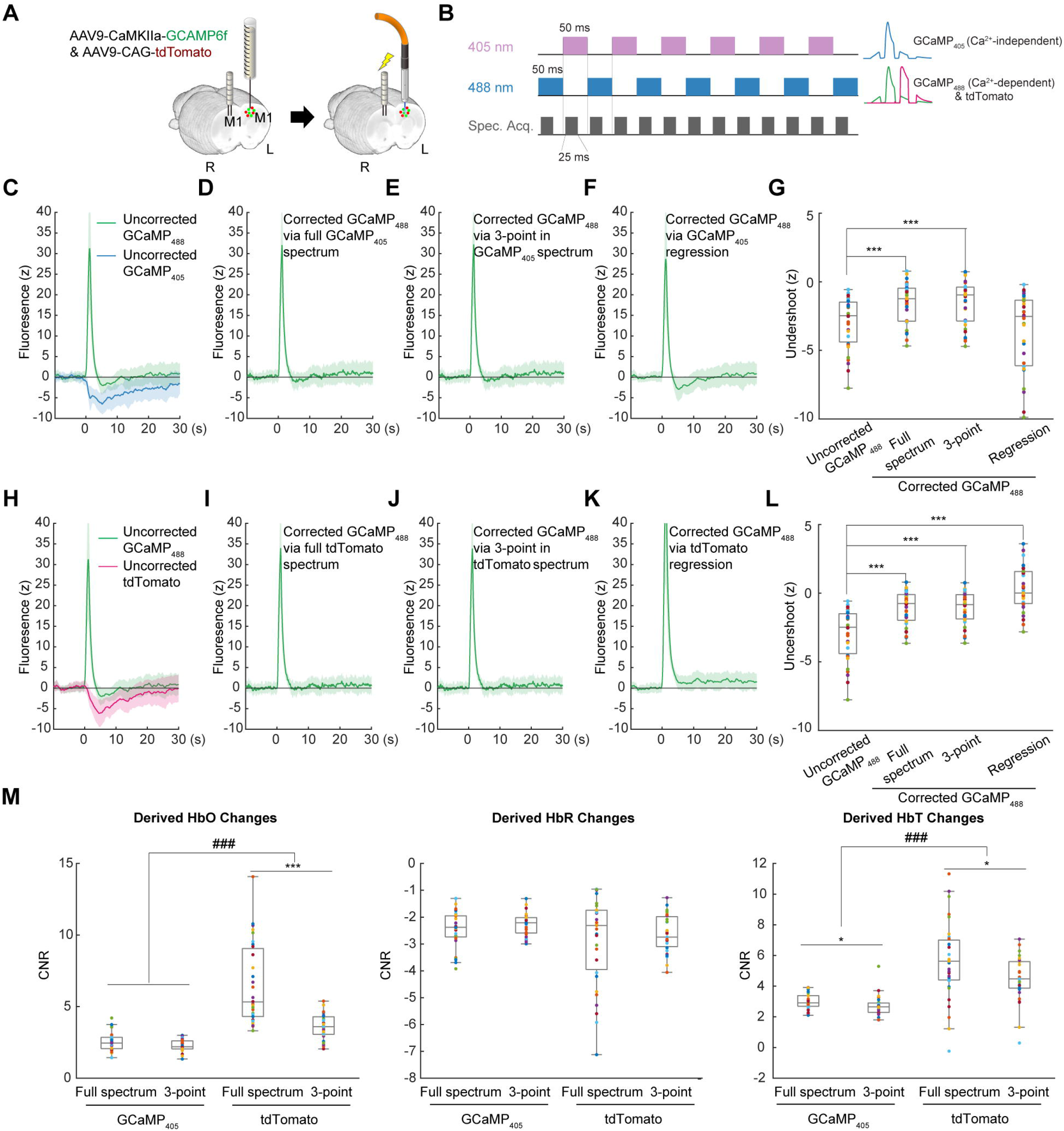
Correction of Hb-absorption artifacts in other fiber-photometry settings. **(A)** Preparation for fiber-photometry measurement of GCaMP and tdTomato signals in the left M1, with the right M1 receiving local micro-electrical stimulation. **(B)** Interleaved acquisition paradigm with 405 nm and 488 nm excitation lasers. Ca^2+^-independent GCaMP signals (GCaMP_405_) were excited by a 405 nm laser, whereas Ca^2+^-dependent GCaMP signals (GCaMP_488_) and tdTomato signals were excited by a 488 nm laser. This interleaved paradigm was used because GCaMP_405_ and GCaMP_488_ share nearly identical emission spectra and thus cannot be spectrally unmixed. Data were acquired at 50% duty cycle every 50 ms using a box-paradigm to avoid spectrometer frame loss during the trigger mode. The final sampling rate is 10 Hz for all signals. **(C)** Measured GCaMP_488_ and GCaMP_405_ signals without any correction. **(D)** Corrected GCaMP_488_ using HbO and HbR dynamics derived from the full GCaMP_405_ spectrum. **(E)** Corrected GCaMP_488_ using HbO and HbR dynamics from 3 datapoints in the GCaMP_405_ spectrum (at 502, 523, and 544 nm). **(F)** Corrected GCaMP_488_ by regressing out the GCaMP_405_ signal dynamics. **(G)** Comparisons of the post-stimulus undershoot artifacts among uncorrected GCaMP_488_ and corrected GCaMP_488_ signals from three different settings as shown respectively in D-F. \*\*\**p* < 0.001, paired *t*-test (n=31). **(H)** Measured GCaMP_488_ and tdTomato signals without any correction. **(I)** Corrected GCaMP_488_ using HbO and HbR dynamics derived from full the tdTomato spectrum. **(J)** Corrected GCaMP_488_ using HbO and HbR dynamics from 3 datapoints in the tdTomato spectrum (at 578, 594, and 610 nm). **(K)** Corrected GCaMP_488_ by regressing out the tdTomato signal dynamics. **(L)** Comparisons of the post-stimulus undershoot artifacts among uncorrected GCaMP_488_ and corrected GCaMP_488_ signals from three different settings as shown respectively in I-K. \*\*\**p* < 0.001, paired *t*-test (n=31). **(M)** Sensitivity of the HbO, HbR, and HbT derived from full or 3-point in emission spectra. Comparison was made between these two conditions and also between the use of GCaMP_405_ and tdTomato signals. *** *p* < 0.001, paired *t*-test (n=31); ### *p* < 0.001, two-way ANOVA (n=31).

### Brain regional differences in hemodynamic response function

Because our proposed technique allows simultaneous measurements of neuronal activity (e.g., GCaMP) and vascular responses (e.g., Δ*C_HbT_*) with minimized Hb-absorption artifacts, it offers a significant added benefit towards studying the relationship between the two. This relationship, often termed the hemodynamic response function (HRF), governs the data interpretation of many noninvasive neuroimaging technologies such as fMRI, intrinsic optics, or functional ultrasound (Taylor et al., 2018). While the use of one canonical HRF to model activity throughout the brain remains the most common practice in the field, accumulating evidence shows that HRF is region-dependent (Devonshire et al., 2012; Taylor et al., 2018) and that a thorough understanding of HRF affected by regional cell types, neurochemicals, and subject conditions is crucial to accurately interpret most neuroimaging data(Lecrux and Hamel, 2016). To demonstrate the utility of our proposed technique for computing region- and sensor-dependent HRFs (Chao et al., 2022), we first described the method to compute HRF (**Figure S13A**) similar to that in (Chao et al., 2022) and then validated the results on a dataset not used in HRF computation (**Figure S13B**). Next, we virally expressed GCaMP6f using AAV under the CaMKIIα promoter and tdTomato under the CAG promoter in the primary motor cortex (M1) of the left hemisphere and the caudate putamen (CPu, a.k.a. dorsal striatum) of the right hemisphere (**Figure 6A, B**). We implanted a microelectrode into the M1 of the right hemisphere (**Figure 6A**) such that transcallosal and corticostriatal stimulation could be achieved simultaneously. Upon electrical stimulation (60 Hz, 300 μA, 0.5 ms pulse width for 2 s), we observed robust GCaMP changes in the left M1 and right CPu (**Figure 6C, D**), which are both monosynaptically connected to the stimulation target. Following the calculation of Hb concentration changes, Hb-absorption correction, and computation of HRFs, we observed that M1 and CPu exhibited distinct HRF profiles (**Figure 6C, D**). Interestingly, we also observed distinct HbR polarities between these two regions.

**Figure 6.**
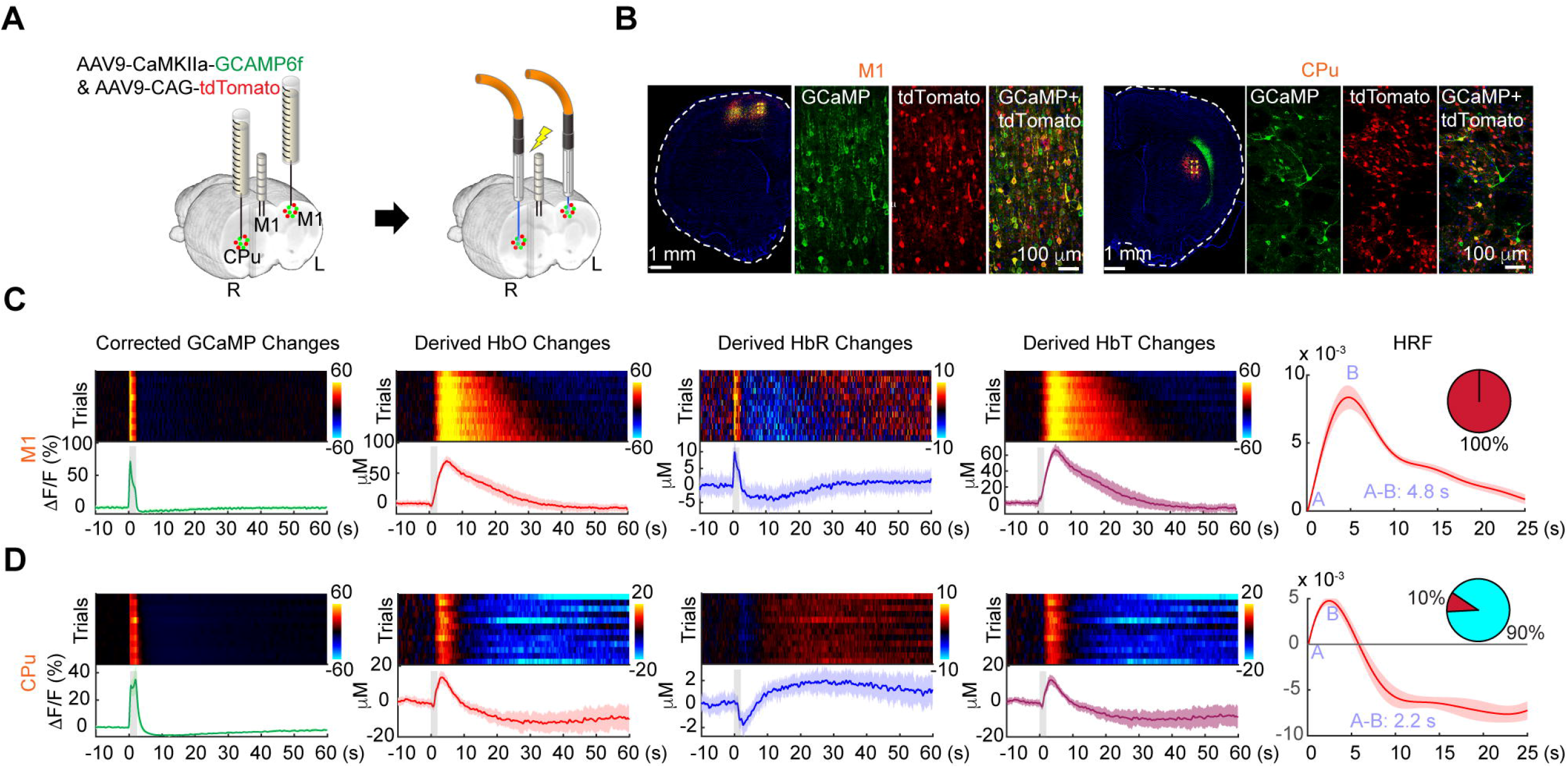
Brain regional differences in hemodynamic response function. **(A)** Preparation for fiber-photometry measurement of GCaMP and tdTomato signals in the left M1 and right CPu, both have direct anatomical connection with the right M1 receiving local micro-electrical stimulation. **(B)** Confocal images showing GCaMP and tdTomato expression in M1 and CPu. (**C, D**) Peri-stimulus heatmaps and response time-courses of the changes in corrected GCaMP, derived HbO, derived HbR, and derived HbT, and the modeled HRFs in M1 and CPu, respectively. Inserted piecharts indicate the ratio between positive (red) and negative (blue) HRF polarities. All per-stimulus heatmap color bars use the same unit as the time-course Y axis. Error bars represent standard deviation. See also Figure S11.

## Discussion

Techniques measuring fluorescence signals in vivo, such as fiber-photometry (Liang et al., 2017; Schlegel et al., 2018; Chao et al., 2022), one-photon microscopy (Bollimunta et al., 2021), two-photon microscopy (Baraghis et al., 2011), or macroscopic imaging through a thinned-skull or implanted-window (Cramer et al., 2021), may all suffer from Hb-absorption. Unlike many imaging techniques that provide spatial information, fiber-photometry does not resolve fluorescence signals emitted from individual cells and its analysis cannot spatially circumvent around the blood vessels. Instead, fiber-photometry utilizes an optical fiber to collect all photons from the targeted volume (**Figure S4A-C**). With the conventional fiber-photometry setting, the targeted volume was reported to be 10^5^ – 10^6^ μm^3^ with a recording depth approximately at 200 μm from the fiber tip (Pisanello et al., 2019). Our simulated detection volume is well within this range (**Figure S14**). With the average blood volume fraction in the brain and Hb concentration documented in the literature (van Zijl et al., 1998), about 9860 μM Hb would appear within the photometry-detectable brain volume and affect the accuracy of fluorescent sensor measurement by absorbing both the excitation and emission photons. In a normal physiological condition, *C_HbO_*, *C_HbR_*, and *C_HbT_* could change for up to 40 μM (Hillman, 2014) and CBV could change for up to ^~^20% (Uludaĝ and Blinder, 2018; Hua et al., 2019) in response to neuronal activity, making such correction to be a time-dependent problem. Our derived changes in Δ*C_HbO_*, Δ*C_HbR_* and Δ*C_HbT_* (**Figure 2F, G, J, Figure 3G-I**), and measured rhoCBV (**Figure 2H**), MRI-CBV (**Figure 3L**) indeed support the amplitude of those changes are possible *in vivo*. Our data indicated that roughly 20% of CBV increase would contribute to 4.08% decrease of green fluorescence signal changes (**Figure S15A**, 95% CI 1.33 - 6.83%), and 2.20% decrease of red fluorescence signal changes (**Figure S15B**, 95% CI 0.57 - 3.84%). Such level of ΔF/F changes on both the green and red sensors are commonly reported in the photometry literature (Chen et al., 2013; Sun et al., 2018) and could be interpreted as the specific sensor activity changes when caution was not exercised. Conventionally, in the brain regions where CBV increases following local neuronal activation, a pseudo-negative fluorescent sensor activity would be expected. However, in cases where CBV decreases with neuronal inhibition (e.g., **Figure 4**), or on occasion with activation (Shih et al., 2009, 2011, 2014; Schridde et al., 2008; Mishra et al., 2011), a pseudo-positive fluorescence signal would occur. Further, while several techniques are readily available to measure CBV (Kwong et al., 1992; Mintun et al., 1984; Sakai et al., 1985; Siegel et al., 2003) (**Figure 2D, H, Figure 3L**), the correction of Hb-absorption cannot be achieved by simply regressing the CBV signal measured concurrently (**Figure S16**). This is mainly because the absorption of HbO and HbR are wavelength dependent and their respective concentrations change over time (Meng and Alayash, 2017; Prahl, 1999) (**Figure S1**). Our simulation suggested that the accuracy of fluorescence signal could be compromised even without a net CBV change (see **Figure S4D** for plots without shaded background). This makes it crucial to quantify both Δ*C_HbO_* and Δ*C_HbR_*, so that the true fluorescence signal changes can be restored.

At present, two major fluorescence-measurement strategies are used in fiber-photometry. One is non-spectrally resolved using a narrow bandpass filter paired with a photodetector (Gunaydin et al., 2014; Kim et al., 2016; Lazarjan et al., 2021). The other is spectrally resolved using a photomultiplier tube array (Cui et al., 2014, 2013) or a spectrometer (Meng et al., 2018; Sun et al., 2018; Chao et al., 2022). The non-spectrally resolved method has the advantage of recording fluorescence signals at a faster speed and can be used with modulated excitation lights and a lock-in amplifier (Gunaydin et al., 2014). In contrast, the spectrally resolved method provides wavelength-dependent information for the fluorescence where the detected photons can be matched with the specific spectral profile of the sensor to improve specificity, or perform spectral unmixing for multiplexing (Meng et al., 2018). If a single nonspectral photodetector is used and Δ*C_HbO_* and Δ*C_HbR_* cannot be obtained, the Hb-absorption spectra (**Figure S1**) and our simulation results (**Figure S4D**) indicated that far-red sensors are less prone to Hb-absorption artifacts. This highlights the importance of continually developing fluorescent sensors in the further red-shifted and far-red range (Patriarchi et al., 2020; Shcherbakova, 2021).

While we did not obtain photodetector-based data in this work, we mimicked the photodetector-based setting by sampling data only from three specific spectral datapoints to enter our Hb-absorption correction pipeline. These modeling data support the utility of our proposed technique in the photodetector-setting (Figure 5E & J). Of note, the detector inside the spectrometer used in this study is a high-end back-thinned CCD array (90% quantum efficiency) with TE cooling (40-50 degrees below ambient) to reduce the dark current (Ocean Insight, 2022). For a rough comparison of the reported detector sensitivity, the most common commercial photodiode detectors used in the fiber-photometry have a sensitivity (reported as responsivity) of 0.38 A/W at 550 nm (Doric, 2021), which is similar to 0.40 A/W at 550 nm of the system used in this study (calculated based on 90% quantum efficiency). It should also be noted that we do not have access to ground-truth GCaMP signals (i.e., the “real” GCaMP signal) using fiber-photometry to make an unequivocal comparison. However, GCaMP measurements from cultured neurons without Hb or using two-photon imaging to circumvent blood vessels have collectively demonstrated the absence of post-stimulus undershoot (Chen et al., 2013; Mittmann et al., 2011). Leveraging these observations in the literature, together with the results we obtained in **Figures 2E and 3F**, we believe in the utility of our proposed Hb-absorption correction method. We recognize that the post-stimulus undershoot artifacts may not significantly compromise all fiber-photometry studies, especially for those studying brief time-locked stimulations/tasks. Nevertheless, we consider the post-stimulus undershoot artifact to be a useful marker to benchmark methods dealing with Hb-absorption and thus used it throughout our studies. We suggest caution be exercised when studying spontaneous ongoing neuronal/neurochemical activity dynamics or slower brain state changes using fiber-photometry because the HbO and HbR concentration changes may cause erroneous readings of the desired sensory activity (see **Figures 4, S11, and S12**). Our results also suggested that regression-based model may be suboptimal to restore the GCaMP signal changes in fiber-photometry (**Figures 5F and K**), and highlighted the benefits in CNR when using a spectrometer system to derive HbO and HbR changes (**Figure 5M**).

The correction method utilizing GCaMP isosbestic excitation (**Figure 5**) is built on the fact that HbO and HbR both present significant absorption at the GCaMP isosbestic excitation wavelength as well as the corresponding emission wavelength. As a result, the fluorescence signal generated from GCaMP isosbestic excitation can be used as the activity-independent fluorescent reference signal in the analytical method proposed in this study and derive Δ*C_HbO_* and Δ*C_HbR_* to restore the functional GCaMP signals. The potential limitations with this approach are (1) trade-off in temporal resolution because interleaved excitation is required, (2) relatively low CNR (**Figure 5M**, note that about one-tenth of the emission signal was obtained as compared to the 488 nm excitation), (3) higher susceptibility to the pH-related confounds (Shaner et al., 2005; Barnett et al., 2017), and (4) though GCaMP has an isosbestic point at an amenable wavelength for this method, other functional fluorescent sensors may perform optimally at a different wavelength (Akerboom et al., 2012; Sun et al., 2020).

Meng et al. recently described the use of spectrally resolved fiber-photometry for simultaneous recording of calcium activities from direct and indirect medium spiny neurons in the CPu and found that both cell types coordinately control the dynamics of movement (Meng et al., 2018). The opportunity to examine the interactions between cell types and/or neurotransmitters has become increasingly popular in fiber-photometry. If both the green and red spectrum are utilized to encode fluorescent sensor activities, it may be possible to use the interleaved sequence as shown in **Figure 5**, or spectral signals from the unmixing residuals (i.e., remaining autofluorescence) to achieve hemodynamic correction (**Figure S17**). In our hands, the residual method suffers significantly from CNR issues due to very low residual photon counts. However, this approach may demonstrate its utility with the advent of more sensitive spectrometer system or improved unmixing algorithm. Considering the increasing need of multiplexing, a valuable next step should consider establishing an experimental pipeline to achieve motion and Hb-absorption artifact-free measurement of a green and a red fluorescent sensors (e.g., GCaMP and jRGECO) in freely-moving animals. This may be achieved by incorporating a far-red activity-independent fluorescent sensor for motion correction first (as Hb-absorption is negligible at this range, see **Figure S1**), followed by Hb-absorption correction using data from isosbestic GCaMP excitation as shown in **Figure 5**.

Given the irreplaceable roles of noninvasive hemodynamic-based neuroimaging techniques (such as fMRI, near-infrared, intrinsic optics, functional ultrasound, positron emission tomography, etc.) in studying brain function, identifying brain region-dependent HRF is crucial towards an accurate interpretation of hemodynamic-based neuroimaging data. Notably, the current standard fMRI analysis uses one HRF shape across the entire brain. Accumulating literature has pointed to the potential problems with this analysis pipeline because HRF is brain region-dependent (Devonshire et al., 2012; Ekstrom, 2021; Jin et al., 2020; Sloan et al., 2010). The current study and other pioneering studies (Albers et al., 2017; He et al., 2018; Liang et al., 2017; Schlegel et al., 2018; Schmid et al., 2016; Schulz et al., 2012; Schwalm et al., 2017; M. Wang et al., 2018; Zhang et al., 2022; Chao et al., 2022) have together demonstrated that fiber-photometry could be seamlessly coupled with fMRI and provide ground-truth cellular activity to help unravel the relationship between neuronal and hemodynamic responses. These studies have bridged a crucial knowledge gap in interpreting fMRI data. The method proposed in this study further expands the scope of this research area, as it generates hemodynamic signals “free-of-charge”, with sensitivity better than fMRI (**Figure 3N**), and the results can be readily compared with the concurrently recorded fluorescent sensor activity. With the spectral photometry recording becoming increasingly popular, the proposed method opens up a unique opportunity to compute HRF in a celltype dependent, stimulation-circuit dependent, and neurotransmission-dependent manner. We expect that this information can be obtained by performing meta-analysis on fiber-photometry data shared through open-science platform and contribute significantly toward a more accurate interpretation of hemodynamic-based neuroimaging results.

In this study, we demonstrated that significant Hb-absorption artifacts in fiber-photometry data may cause inaccurate measurements and therefore erroneous interpretations. We proposed a method that addresses this issue by quantifying dynamic HbO and HbR absorption changes using activity-independent fluorescence spectra, and we demonstrated that these additional quantitative measurements were efficient in correcting the artifact and retrieve the desired, unperturbed fluorescence signals in fiber-photometry. This approach simultaneously measures HbO, HbR, and the activity of a selected fluorescent sensor, providing an additional experimental capability to investigate the influence of neuronal or neurochemical activity on cerebral hemodynamics. Through a proof-of-principle study, we provide direct evidence that HRF is brain-region dependent, suggesting that the analysis of hemodynamic-based neuroimaging data should not apply a single HRF across all brain regions.

## Acknowledgements

We thank UNC CAMRI members and Dr. Elizabeth Hillman for their helpful discussions and critiques. We thank Drs. Lindsay Walton and Dom Cerri for editing the manuscript. This work is supported in part by NIH grants (RF1MH117053, RF1NS086085, R01MH126518, R01MH111429, R01NS091236, P60AA011605, U01AA020023, P50HD103573, S10MH124745, and S10OD026796 to Y.Y.I.S.), the Intramural Research Program of the NIH/NIEHS (1ZIAES103310 to G.C.), the Burroughs Wellcome Fund (CASI Award number 5113244 to N.C.P.), and the Arnold and Mabel Beckman Foundation (2021BYI to N.C.P.).

## Author contributions

W.-T.Z., T.-H.H.C., G.C., and Y.-Y.I.S. conceived the project and designed the experiments. W.-T.Z., T.-H.H.C., T.-W. W., E.A.O., J.Z., and R.N. implemented the methods. W.-T.Z., T.-H.H.C, Y.Y., S.-H.L., and N.C.P. analyzed data. W.-T.Z., T.-H.H.C, and Y.-Y.I.S. wrote the manuscript with input from all authors. H.Z., G.C, and Y-Y.I.S. supervised the study and provided funding support.

## STAR★METHODS

### KEY RESOURCES TABLE

| REAGENT OR RESOURCE | SOURCE | IDENTIFIER |
| --- | --- | --- |
| Recombinant DNA |  |  |
| AAV5-CaMKIIα-EYFP | UNC Vector Core | N/A |
| AAV9-CaMKIIα-GCaMP6f-WPRE-SV40 | U.Penn Vector Core | AV-9-PV3435 |
| AAV9-CAG-tdTomato | UNC Vector Core | N/A |
| AAV9-CaMKIIα-Chr2-EYFP | UNC Vector Core | N/A |
| Software and Algorithms |  |  |
| MATLAB | Mathworks | R2019b |
| RStudio | RStudio Inc. | Version 1.0.136 |
| Other |  |  |
| 488 nm laser and remote | Coherent | OBIS 488 LS (60 mW) |
| 561 nm laser and remote | Coherent | OBIS 561 LS (50 mW) |
| Spectrometer | Ocean Optics | QE-pro |
| Optic patch cable | Thorlabs | FT200EMT (200 mm, 0.39 NA) |
| Uncleaved fiber optic cannula | Thorlabs | CFMC12U-20 (200 mm, 0.39 NA) |
| Bipolar Tungsten electrode | P1 Technologies | MS303T/2C-B/SP ELEC .008/.2MM<br>TW TUNGSTEN (MRI) |

### Monte Carlo simulation for photon transport in fiber-photometry recording

The Monte Carlo simulation of photon transport during fiber-photometry recording was achieved in two steps: 1) excitation light path simulation and 2) emission light path simulation. The goal was to first build a database composed of a large number of possible excitation light paths. After which, an emission starting point can be randomly selected from the excitation light path database for emission light path simulation.

For the excitation light path simulation, we built the database by repeatedly launching excitation photon packets that started with a unity weight = 1. The photon packets deposited weight due to absorption at each step of traveling through the simulated brain, and the photon packet’s current locations and remaining weights were logged along the way. Each excitation photon packet repeatedly underwent the following steps (**Figure S3**, left panel), until it was either terminated or transmitted out of the brain.

#### Step 1: Launching a photon packet (Figure S3, left panel a and b)

In our model, we launched our photon packets from an optic fiber tip with a diameter of 200 μm; the range of possible launch directions, or so-called acceptance angles, would be limited by the fiber numerical aperture (NA) according to Snell’s law. For applications, the NA is most commonly expressed as:

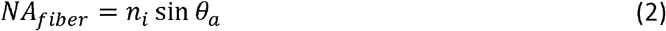

where *θ_a_* is the maximum 1/2 acceptance angle of the fiber, and *n_i_* is the index of refraction of the material outside of the fiber. With this in mind, we simply set the initial launch position randomly within the surface of fiber tip, along with a random initial launch direction that was within the range of acceptance angles. This can be easily done by using three Cartesian coordinates (*x*, *y*, *z*) and three direction cosines (*v_x_*, *v_y_*, *v_z_*) to determine the initial position and direction, respectively.

For initial position:

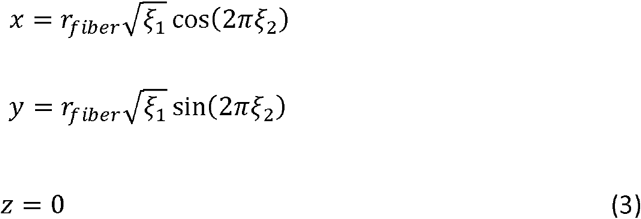

For initial direction:

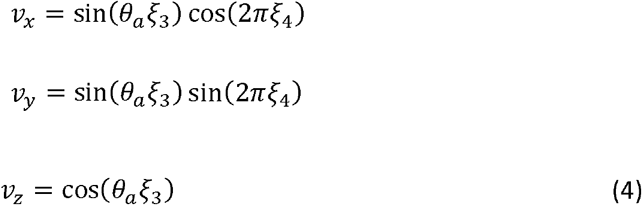

where *r_fiber_* is the radius of the optic fiber, and *ξ*_1–4_ are the independent random numbers between 0 and 1.

#### Step 2: Step size selection and photon packet movement (Figure S3, left panel 1-2 and right panel 5-6)

The step size s is the distance that the photon packet travels between interaction sites. In our study, we selected a step size using the following function derived from the inverse distribution method and the Beer–Lambert law:

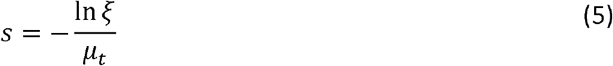

where *ξ* is a random number between 0 and 1, and *μ_t_* is the total interaction coefficient (the sum of the absorption coefficient (*μ_a_*) and scattering coefficient (*μ_s_*) of brain tissue). The *μ_a_* and *μ_s_* for distinct wavelengths of a traveling photon were derived from the optical absorption and scattering properties provided by an online database from Oregon Medical Laser Center, Oregon Health & Science University (https://omlc.org/software/mc/mcxyz/index.html). We assumed 3% blood with 75% average oxygen saturation in our simulated brain.

Once the step size was selected, the photon packet traveled for distance s in the direction defined by direction cosines, and the coordinates were updated as follows:

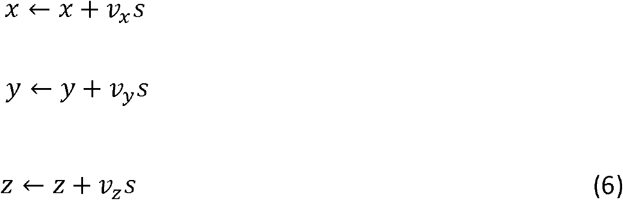

#### Step 3: Absorption and scattering (Figure S3, left panel 3-4 and right panel 7-8)

At each interaction site, photon weight was partially absorbed by brain tissue, and the decreased fraction of the weight Δ*w* can be defined as:

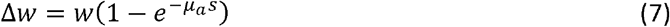

The weight of the photon packet was then updated as follows:

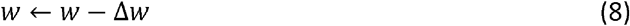

Following absorption, the photon packet was scattered. The scattering direction was defined by both scattering angle *θ* and polar angle *φ*. The scattering angle was determined using the Henyey-Greenstein phase function(Henyey and Greenstein, 1941):

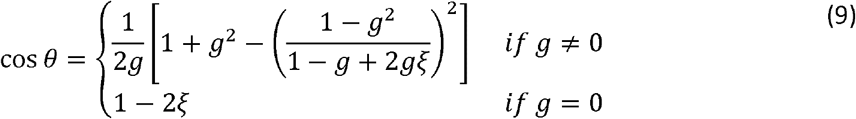

where *g* refers to scattering anisotropy, which has a value between −1 and 1. When *g* is 0, it generally indicates that the scattering is isotropic. As *g* approaches 1 or −1, the scattering is confined to forwards and backwards, respectively.

The polar angle was assumed to be uniformly distributed between 0 and 2π, as below:

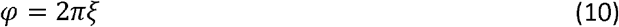

With these angles and the original direction cosines, we were able to calculate the new set of propagation direction cosines, which can be represented in the global coordinate system as follows:

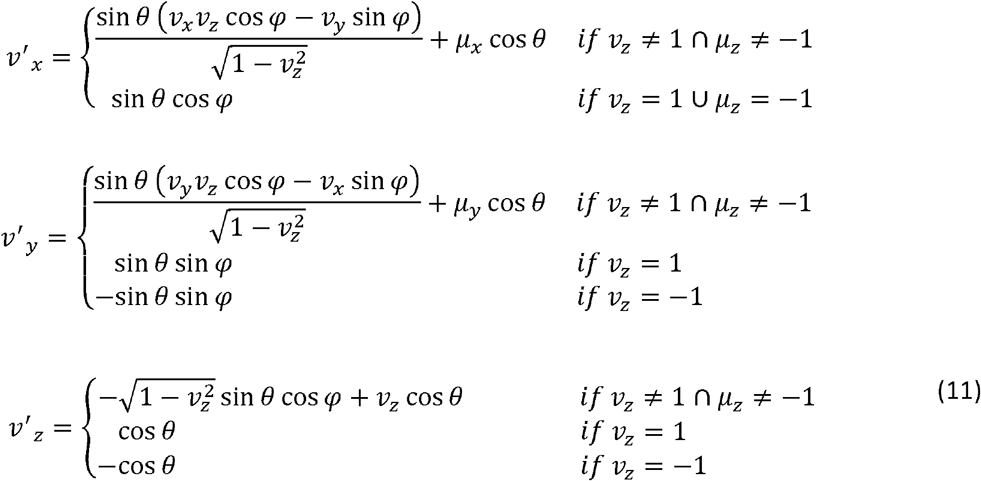

#### Step 4: Photon termination

Steps 2 and 3 were repeated until the weight of the photon packet reached 10^-4^ or less, then the simulation for this particular photon packet was terminated. The excitation light path and weight of each step were saved to the database, after which a new photon packet simulation was started from step 1.

The emission light path simulation was similar to the excitation light path simulation. Specifically, we need to first determine the initial position, the initial direction, and the initial weight before launching an emission photon packet (**Figure S3**, right panel c-d). The initial coordinates of each emission photon packet were randomly selected from the excitation light path database. All coordinates in the database had an equal probability to be selected regardless of excitation light path. If the selected coordinates were within an area containing fluorophore, its initial coordinates were determined first, and then the corresponding excitation light path history (from the start to the point where the emission occurred) was recorded. In this study, we assumed the fluorophore existed equally within a 1 mm diameter sphere of viral expression area centered 0.3 mm below the fiber tip.

The initial direction of an emission photon was assumed to have isotropic probability in all directions; therefore, the direction cosines can be derived using the following:

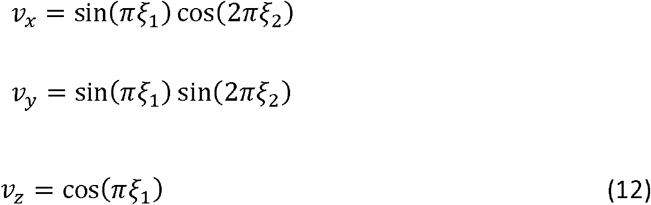

Lastly, the initial weights were inherited from the remaining weights of the excitation photon packet at the initial coordinates.

Steps 2 and 3 were repeated until the weight of the photon packet reached 10^-4^ or less, and then the simulation of this photon packet was terminated (step 4). In emission step 4, only when the photon packet reached the fiber tip at smaller than 1/2 acceptance angle and a weight beyond 10^-4^, the paired excitation/emission light path and the weight of each step were saved to the database. In order to improve signal-to-noise ratio, we collected 2-2.5 × 10^4^ pairs of excitation and emission light paths.

The final weight (*W_final_*) when the emission photon reaching the optical fiber tip represents the survival rate of the current combined excitation/emission light path. Therefore, the total average pathlength of excitation and emission lights are then calculated by summing each survived photon’s *W_final_* multiplied by its paired excitation and emission pathlength respectively:

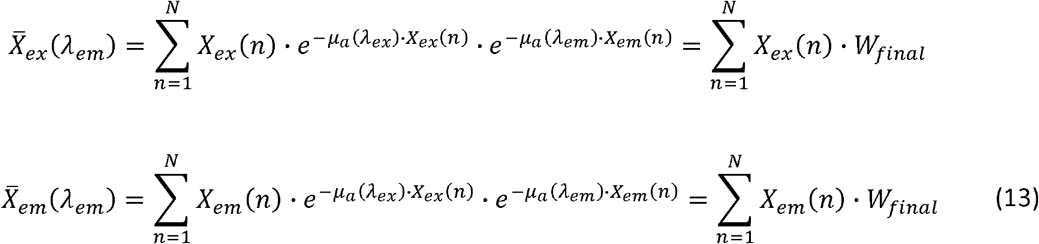

Since the *W_final_* is affected during both excitation and emission phase, the emission measured at different wavelengths have not only distinct average emission pathlength 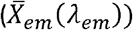, but also distinct average excitation pathlength 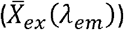 even when these emission wavelengths are excited by the same excitation wavelength. In this study, only the 488 nm excitation wavelength was used and simulated.

To plot the 488 nm excitation light illuminating distribution (Figure S 3a), all excitation light paths’ coordinates generated from the excitation light path simulation, regardless to the emission light detection, were first compressed into 2D coordinates, and then used for the 2D histogram. To plot the 515 or 580 nm emission light receptive probability map (Figure S 3b-c), the initial emission 3D coordinate of each successfully detected 515 or 580 emission photons were compressed into 2D coordinates and shown in a 2D histogram.

### Correction of Hb-absorption

Given the different absorption of HbO and HbR (**Figure S1**), their dynamic concentration changes must be quantified in order to properly correct fluorescent sensor signals. To address this, we proposed the following methods to derive HbO and HbR concentration (Δ*C_HbO_* and Δ*C_HbR_*) time-courses using spectral datapoints.

In a simple, non-absorbing and non-scattering medium, the measured fluorescence signal *F* at any time *t* is linearly related to the fluorescent protein concentration *C*(*t*):

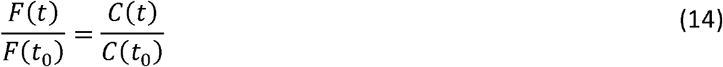

where *F*(*t*_0_) and *C*(*t*_0_) represent the fluorescence signal and the fluorescent protein concentration at the reference time point *0*, respectively. When the absorption of the medium is considered, and *F* is measured at multiple wavelengths *λ_Em_*, equation (14) can be further modified according to an established model (Ma et al., 2016):

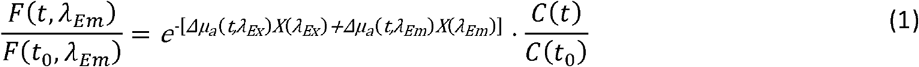

Then after natural logarithmic transformation:

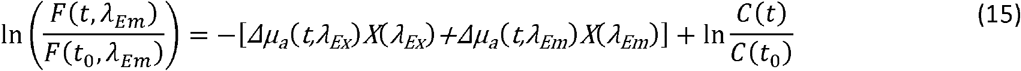

where *Δμ_a_*(*t,λ_Ex_*) represents the change of the absorption coefficient *μ_a_* at excitation wavelength *λ_Ex_* at the time point *t* compared to the time point *0; Δμ_a_*(*t,λ_Em_*) represents the change of the absorption coefficient *μ_a_* at emission wavelength *λ_Em_* at the time point *t* compared to the time point *0*; *X*(*λ_Ex_*) and *X*(*λ_Em_*) represent the photon traveling pathlengths at the given wavelength, respectively. Next, HbO and HbR concentration changes (Δ*C_HbO_* and Δ*C_HbR_*) as a function of time need to be incorporated. When using 488 nm excitation, the absorption term in equation (15) that contains *Δμ_a_* could be expressed as:

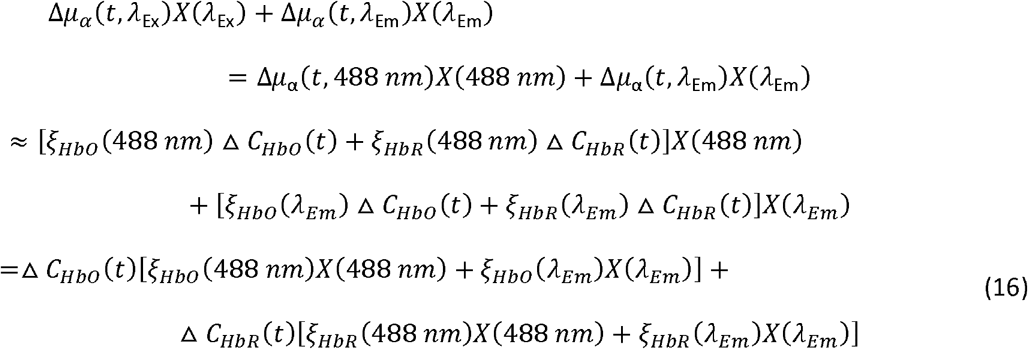

where *ξ_HbO_* and *ξ_HbR_* are well-established molar extinction coefficients at different wavelengths of HbO and HbR, respectively (Prahl, 1999).

At the time point *t*, the measured fluorescence ratio after natural logarithmic transformation, ranged from *λ*_*Em*_1__ to *λ_Em_n__* could be put together in a vector form *M*(*t*):

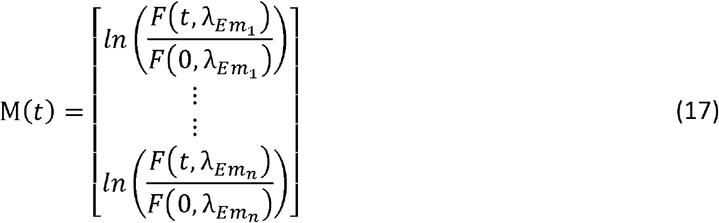

Additionally, two vectors A and B could be used to represent the following:

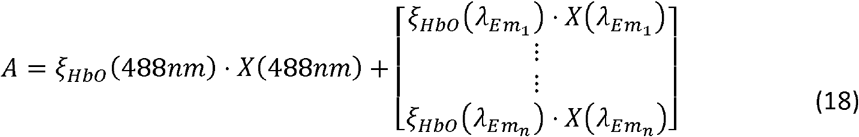

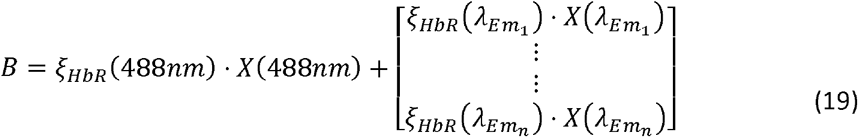

With any activity-independent fluorescence signals, the last component 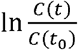 should be 0. Practically, we designate an error value σ. Taken together, equation (15) could then be simplified and reorganized as equation (20). To solve Δ*C_HbO_*(*t*) and Δ*C_HbR_*(*t*) at any time point *t*, we used generalized method of moment (GMM)(Hansen, 1982) to minimize L in equation (21). By performing the GMM computation for each time point, we can obtain time-courses of Δ*C_HbO_*(*t*), Δ*C_HbR_*(*t*) and *σ*.

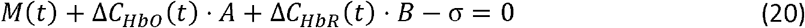

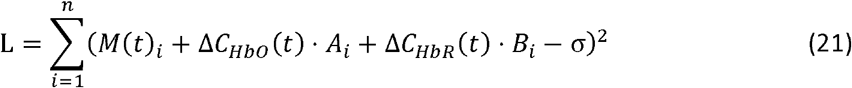

In this case, we have numerous spectral datapoints that can be used to resolve two unknowns that are of interest – Δ*C_HhO_*(*t*) and Δ*C_HbR_*(*t*). After which, Δ*μ_α_*(*t,Λ*_Ex_) and Δ*μ_α_*(*t, λ*_Em_) can be calculated using equation (16) and *F*(*t, λ_Em_*) can be subsequently calculated using equation (15) to correct for HbO and HbR absorptions.

### Subject

This study employed a total of 26 male Sprague Dawley (SD) rats weighing between 300-600g. All procedures were performed in accordance with the National Institutes of Health Guidelines for Animal Research (Guide for the Care and Use of Laboratory Animals) and approved by the University of North Carolina (UNC) Institutional Animal Care and Use Committee. These rats were separated into 6 cohorts. In the first cohort (n=3), EYFP was expressed in the bilateral S1FL using AAV5-CaMKIIα-EYFP (titer 4.2 × 10^12^ molecules/mL, UNC Vector Core) to demonstrate the impact of Hb concentration changes on activity-independent fluorescence signals and evaluate the proposed correction method. In the second cohort (n=4), GCaMP and tdTomato were co-expressed in the bilateral S1FL using a mixture of AAV9-CaMKIIα-GCaMP6f-WPRE-SV40 (titer ≥ 1 × 10^13^ vg/mL, Penn Vector Core) and AAV9-CAG-tdTomato (titer 3.8 × 10^12^ molecules/mL, UNC Vector Core) at 5:1 ratio to demonstrate the impact of Hb concentration changes on a commonly used fluorescent sensor and evaluate the proposed correction method. In the third cohort (n=3), GCaMP and tdTomato were co-expressed in the left PrL using a viral mixture identical to the second cohort. The fourth cohort (n=4) received a twisted tungsten electrode implanted into the M1, and GCaMP and tdTomato were co-expressed in the contralateral M1 using a viral mixture identical to the second cohort. The fifth cohort (n=5) was used in a supplementary study of isoflurane challenge, and only had GCaMP expressed in the left M1. The sixth cohort (n=4) received a twisted tungsten electrode implanted into the M1, and GCaMP and tdTomato were co-expressed in the contralateral M1 and ipsilateral striatum using a viral mixture identical to the second cohort. The seventh cohort (n=3) was used in a supplementary study, and had channelrhodopsin2 (ChR_2_) expressed in the ventroposterior thalamus (VP) using AAV9-CaMKIIα-ChR2-EYFP (titer 3.6 X 10^12^ molecules/mL, UNC Vector Core) and GCaMP expressed in the ipsilateral S1 using AAV9-CaMKII-GCaMP6f-WPRE-SV40 (titer ≥ 1 × 10^13^ vg/mL, Penn Vector Core). All rats in this study were housed under environmentally-controlled conditions (12 h normal light/dark cycles, lights on at 7am; 20–23 °C and 40–60% relative humidity), with ad libitum access to food and water.

### Stereotactic surgery

For all surgical procedures, rats were anesthetized initially by 5% isoflurane, and maintained by a constant flow of 2-3% isoflurane mixed with medical air. Rectal temperature was continuously monitored and maintained within 37□±□0.5 °C using a feedback-controlled heating pad (Harvard Apparatus, Model 557020, Holliston, MA). Rats were head-fixed to a stereotactic frame (Kopf Instruments, Model 962, Tujunga, CA). The skin was opened to expose the skull surface, and burr holes were prepared according to experimental coordinates. All coordinates used for microinjection are listed as follows. S1: AP = +0.5 mm and ML = +3.7 mm, DV = 1.3 mm, M1: AP = +1.8 mm and ML = −2.5 mm, DV = 2.0 mm, Striatum: AP = 0.5 mm and ML = 4.0 mm, DV = 5.2 mm, VP: AP = −2.8 mm and ML = +2.5 mm, DV = 6.5 mm, PrL: AP = 2.2 mm and ML = 0.9 mm, DV = 4.1 mm. Microinjections were performed at a flow rate of 0.1□μL/min for 1 μL, and an additional 10□min was given for virus diffusion prior to slow retraction of the microsyringe needle. The burr holes were then sealed with bone wax (Fisher Scientific, Pittsburgh, PA), and the wound was sutured. A month after the virus microinjection, the skin was reopened to expose the skull, then the bone wax was removed and optical fibers (200□μm in diameter; NA: 0.39) were chronically implanted to coordinates 0.2-0.5 mm above the virus injection sites. Four MR-compatible miniature brass screws (Item #94070A031, McMaster Carr, Atlanta, GA) were anchored to the skull, then the surface of the skull was covered with dental cement to seal implanted components and the wound was sutured to further protect the surgical site.

At the end of every surgical procedure, lidocaine jelly (#866096, Henry Schein Inc., Melville, NY) was applied around the surgical wound for pain relief, and to prevent the rat scratching the wound. Meloxicam (#6451720670, Henry Schein Inc., Melville, NY) was also given by oral administration for further pain relief. Rats were allowed at least 1 week for recovery from surgical procedures before any further experiments.

### Experimental setup

The spectrally resolved fiber-photometry system in this study replicates an established system described previously(Meng et al., 2018). Laser beams from a 488 nm 60 mW continuous wave (CW) laser (OBIS 488 LS-60, Coherent, Santa Clara, CA) and a 561 nm 50 mW CW laser (OBIS 561 LS-50, Coherent, Inc.) are aligned and combined by broadband dielectric mirrors (BB1-E02, Thorlabs, Newton, NJ) and a long-pass dichroic mirror (ZT488rdc, Chroma Technology Corp), then launched into a fluorescence cube (DFM1, Thorlabs, Newton, NJ). Extra neutral density filters (NEK01, Thorlabs, Newton, NJ) are placed between the combined laser beam and the fluorescence cube to adjust the final laser power. The fluorescence cube contains a dichroic mirror (ZT488/561rpc, Chroma Technology Corp) to reflect and launch the combined laser beam through an achromatic fiber port (PAFA-X-4-A, Thorlabs, Newton, NJ) into the core of a 105/125 mm core/cladding multi-mode optical fiber patch cable (FG105UCA, Thorlabs, Newton, NJ). The distal end of the patch cable is connected to an implantable optical fiber probe for both excitation laser delivery and emission fluorescence collection. The emission fluorescence collected from the fiber travels back along the patch cable into the fluorescence cube, passes through the dichroic mirror and an emission filter (ZET488/561 m, Chroma Technology Corp, Bellows Falls, VT), then launches through an aspheric fiber port (PAF-SMA-11-A, Thorlabs, Newton, NJ) into the core of an AR-coated 200/230 mm core/cladding multi-mode patch cable (M200L02S-A, Thorlabs, Newton, NJ). The AR-coated multi-mode patch cable is connected to a spectrometer (QE Pro-FL, Ocean Optics, Largo, FL) for spectral data acquisition, which can be operated by a UI software OceanView (Ocean Optics, Largo, FL). In order to achieve concurrent recording during fMRI, trigger mode is used in OceanView, where the photometry system is synchronized with MRI using an Arduino micro-controller board.

The stimulation system uses a DAQ board (1208Hs-2AO, Measurement Computing Corp., Norton, MA) to send out stimulus triggers according to the stimulus paradigm set in a homemade software program. During fMRI experiments, the DAQ synchronizes stimulation pulses via triggers from the MRI system. Stimulation pulses were driven by a constant current stimulus isolator (A385RC, World Precision Instruments, Sarasota, FL) for forepaw and microelectrical stimulation experiments; and by a 473 nm blue diode laser (Shang Laser and Optics Century, BL473T8-200FC + ADR-800A, Shanghai, China) for optogenetic experiments.

### Animal subject preparation and physiology management

Rats were initially anesthetized with 4% isoflurane (Vaporizer #911103, VetEquip Inc., Livermore, CA, USA) mixed with medical air and endotracheally intubated using a 14G x 2”(>400 g) or 16G x 2”(<400 g) i.v. catheter (Surflash Polyurethane Catheter, TERUMO, Somerset, NJ, USA). Respiration was maintained by a ventilator (SAR-830 or MRI-1, CWE Inc, Ardmore, PA, USA) set at 60 breaths/min and an inspiration time ratio of 40%. A rectal probe was used to monitor core body temperature (OAKTON Temp9500, Cole-Parmer, Vernon Hills, IL, USA) and a capnometer was used to monitor heart rate, peripheral blood oxygen saturation, and end-tidal CO_2_ (SURGIVET^®^ V90041LF, Smith Medical, Dublin, OH, USA). Body temperature was maintained at 37 ± 0.5°C using a circulating water blanket connected to a temperature adjustable water bath (Haake S13, Thermo Fisher Scientific, Waltham, MA, USA). Ventilation tidal volume was adjusted to keep the heart rate at 300 ± 50 beats per minute, peripheral blood oxygen saturation above 90%, and end-tidal CO_2_ between 2.8 - 3.2%. End-tidal CO_2_ values from this capnometer system were previously calibrated against invasive sampling of arterial blood gas, reflecting a partial pressure of carbon dioxide (pCO_2_) level of 30–40 mm Hg(Shih et al., 2012, 2013). For studies using Rhodamine B for CBV measurements, a bolus dose of 40 mg/kg (Sigma–Aldrich, St. Louis, MO) was injected via tail vein.

### Concurrent functional MRI Scan with fiber-photometry recording

All fMRI data in this study were collected on a Bruker BioSpec 9.4-Tesla, 30 cm bore system with 6.0.1 on an AVANCE II console (Bruker BioSpin Corp., Billerica, MA). An RRI BFG 150/90 gradient insert (Resonance Research, Inc, Billerica, MA) paired with a Copley C700 gradient amplifier (Copley Controls Corp., Canton, MA) was used. A homemade single-loop surface coil with an internal diameter of 1.6□cm was used as a radio-frequency transceiver. Isoflurane concentrations were adjusted to 2% and animals were secured in to a custom-built, MR-compatible rat cradle. Animal physiology was monitored and maintained as described in the previous paragraph.

Upon stabilizing the animals, a pair of needle electrodes was inserted under the skin of forepaw for stimulation, or electrode/optical fiber patch cables were connected according to experimental design. Before connecting the fiber-photometry patch cable, all light in the room was turned off, the final output power of the 488 nm laser and 561 nm laser were 51.3 ± 18.6 μW (mean ± SD, n = 26), which were adjusted to balance spectral amplitudes (Meng et al., 2018). 100 μW power is the maximum we suggested when the virus expression is less robust. This power range is consistent with published fiber-photometry studies (Gunaydin et al., 2014; Jones-Tabah et al., 2021, p.; Liang et al., 2017; Zhang et al., 2021). A background spectrum was measured as a reference by pointing the fiber tip to a nonreflective background in the dark room. This background spectrum was then automatically subtracted by OceanView during photometry recording.

Following setup processes, the cradle was pushed into MRI bore, and a bolus of dexmedetomidine (0.025 mg/kg; Dexdormitor, Orion, Espoo, Finland) cocktailed with paralytic agent rocuronium bromide (4.5 mg/kg; Sigma–Aldrich, St. Louis, MO) was injected into the tail vein (Chao et al., 2018). Fifteen minutes after the bolus injection, continuous intravenous infusion of dexmedetomidine (0.05 mg/kg/h) and rocuronium bromide (9 mg/kg/h) cocktail was initiated and the isoflurane concentration was adjusted to 0.5–1% for the entire scanning period.

Magnetic field homogeneity was optimized first by global shim and followed by local first- and second-order shims according to B0 map. Anatomical images for referencing were acquired using a rapid acquisition with relaxation enhancement (RARE) sequence (12 coronal slices, thickness = 1 mm, repetition time (TR) = 2500 ms, echo time (TE) = 33 ms, matrix size = 256 × 256, field-of-view (FOV) = 25.6 × 25.6 mm^2^, in plane resolution 0.1 x 0.1 mm, average = 8, RARE factor = 8). The center of the 5^th^ slice from the anterior direction was aligned with the anterior commissure. Cerebral blood volume (CBV) fMRI scans were acquired using a multi-slice single-shot gradient echo echo-planar imaging sequence (GE-EPI) (slice thickness□=□1□mm, TR□=□1000□ms, TE□=□8.1□ms, matrix size□=□80□×□80, FOV□=□25.6□×□25.6□mm^2^, in plane resolution 0.32 x 0.32 mm, bandwidth□=□250□kHz), with the same image slice geometry imported from the previously acquired T2-weighted anatomical image. A session of GE-EPI scans with 300 repetitions was taken, and at about the 100^th^ scan, Feraheme (30□mg Fe/kg, i.v.) was administered for CBV percentage change calculations.

### CBV fMRI Data Processing and Statistical Analyses

All fMRI data were analyzed using the Analysis of Functional Neuroimages (AFNI)(Cox, 1996) general linear model (GLM) framework(Worsley and Friston, 1995). All EPI images were skull-stripped, and slice-timing was corrected. Then automatic co-registration was applied to realign time-courses within subjects to correct subtle drift of EPI images. Finally all EPI images were aligned to a T2-weighted rat brain template(Valdés-Hernández et al., 2011) to generate normalized fMRI images to allow for group-level comparisons. Gaussian smooth (FWHM = 0.5 mm) was performed before feeding into the GLM fitting. To test the group-level significant consistency of the stimulus-evoked responses, we employed a parametric one-sample t-test implemented in AFNI. The false discovery rate correction was used to adjust for the multiple comparisons of fMRI maps (p < 0.05). A 3×3 voxel region of interest (ROI) was placed at the fiber tip in the S1 to extract fMRI time-course data. To account for Feraheme kinetics over the course of experiments, the following equations were used to calculate the 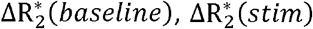, and ΔCBV (Decot et al., 2017).

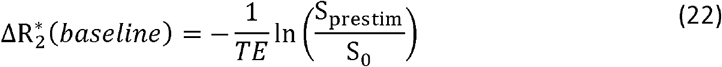

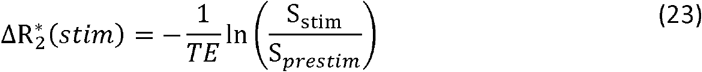

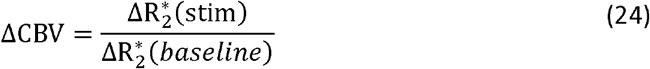

where *S_prestim_* and *S*_0_ represents MR signal intensity after and before Feraheme injection, respectively, and *S_stim_* are the MR signal intensities during the stimulation.

### Fiber-photometry Spectral unmixing

Mixed spectra acquired by fiber-photometry were analyzed using a spectral linear unmixing algorithm(Meng et al., 2018). Briefly, at any time point n, the mixed spectrum *Y*(*n*) was modeled as:

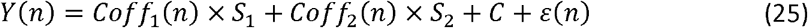

where *S*_1_ and *S*_2_ are the normalized reference emission spectra of the two fluorescence signal sources. Coff_1_ and Coff_2_ are the unknown regression coefficients corresponding to the *S*_1_ and *S*_2_ respectively. *C* is the unknown constant, and *ε*(*n*) is random error. *Coff*_1_(*n*), *Coff*_2_(*n*), and *ε*(*n*) at each time point were estimated using the lm() function in the RStudio package (RStudio Inc. V1.0.136, Boston, MA).

### Modeling the hemodynamic response function

The relationship between neuronal activity and hemodynamic response can be expressed as:

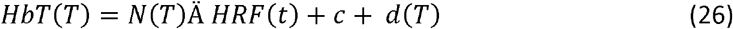

where the *HbT*(*T*) represents hemoglobin fluctuation time-course, the *N*(*T*) represents neuronal activity time-course, the *HRF*(*t*) represents an impulse hemodynamic response function with *t* sampling points, *c* is a constant for baseline offset and *d*(*T*) is for linear drift over time. Assuming *T* = {0, 1, 2,…, *m*}, *t* = {0, 1, 2,…, *n*}, this equation can be expressed as:

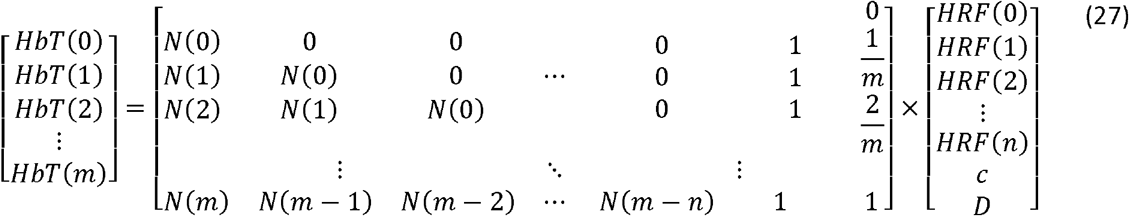

Therefore, the *HRF*(*t*), *c* and *D* (slope of *d*(*T*)) can be solved using Ordinary Least Squares solution with known *HbT*(*T*) and *N*(*T*).

To avoid physiological noise contamination, *HbT*(*T*) were low-pass filtered with a cutoff frequency at 0.5 Hz before the HRF estimation.

### Histology

At the end of the experiments, rats were euthanized by a mixture of 1-2 ml of sodium pentobarbital and phenytoin sodium (Euthasol, Virbac AH, Inc., Westlake, TX), and transcardially perfused with saline followed by 10% formalin. The brains were removed and stored in 10% formalin overnight, then transferred into a 30% sucrose solution (in 0.1 M phosphate buffer) for 2–3 days, until brains sunk to bottom of storage bottles. These brains were cut into serial coronal sections (40 μm) using a cryotome (#HM450, Thermo Fisher Scientific, Waltham, MA) and mounted on glass slides. Fluoro-Gel II Mounting Medium (#17985-50, Electron Microscopy Sciences, Hatfield, PA) was covered on the brain slides to provide DAPI stain and for fluorescence imaging. Slides were imaged using a Zeiss LSM780 confocal microscope.

---

**Table S1:** Average pathlengths *X(λ_Ex_)* and *X(λ_Em_)* by Monte Carlo Simulation.

| Emission wavelength (nm) | Average $X(\lambda_{Ex})$ (cm) | Average $X(\lambda_{Em})$ (cm) |
| --- | --- | --- |
| 488 | 0.045093573 | 0.044381928 |
| 489 | 0.046031195 | 0.045206893 |
| 490 | 0.045353095 | 0.044599308 |
| 491 | 0.045145851 | 0.043833332 |
| 492 | 0.044940114 | 0.044644849 |
| 493 | 0.045780127 | 0.044951492 |
| 494 | 0.044833187 | 0.044238425 |
| 495 | 0.045254011 | 0.044087712 |
| 496 | 0.044832712 | 0.04357339 |
| 497 | 0.045793623 | 0.04438636 |
| 498 | 0.044556411 | 0.043741701 |
| 499 | 0.046485322 | 0.045282627 |
| 500 | 0.044936862 | 0.045399841 |
| 501 | 0.044904189 | 0.043737664 |
| 502 | 0.044676525 | 0.044692953 |
| 503 | 0.046449058 | 0.045104426 |
| 504 | 0.04554069 | 0.045690015 |
| 505 | 0.044186305 | 0.042754367 |
| 506 | 0.044463103 | 0.043549104 |
| 507 | 0.045240076 | 0.04375625 |
| 508 | 0.045271499 | 0.043551134 |
| 509 | 0.045203535 | 0.044181218 |
| 510 | 0.04544563 | 0.04325581 |
| 511 | 0.045264121 | 0.043061521 |
| 512 | 0.044537346 | 0.042787669 |
| 513 | 0.044684362 | 0.042203088 |
| 514 | 0.044171835 | 0.041233961 |
| 515 | 0.044738093 | 0.04226692 |
| 516 | 0.044465855 | 0.041780618 |
| 517 | 0.044712008 | 0.041028167 |
| 518 | 0.043706904 | 0.039055061 |
| 519 | 0.045479041 | 0.040324911 |
| 520 | 0.04389417 | 0.039274782 |
| 521 | 0.043549373 | 0.038231377 |
| 522 | 0.043219292 | 0.037524461 |
| 523 | 0.043155712 | 0.037460255 |
| 524 | 0.042766927 | 0.03694169 |
| 525 | 0.043084675 | 0.035815527 |
| 526 | 0.043150433 | 0.034397857 |
| 527 | 0.043739313 | 0.034308684 |
| 528 | 0.04204147 | 0.033144384 |
| 529 | 0.042283369 | 0.033431559 |
| 530 | 0.041576273 | 0.03274051 |
| 531 | 0.041403969 | 0.032249133 |
| 532 | 0.042259357 | 0.03207917 |
| 533 | 0.042053695 | 0.03107138 |
| 534 | 0.041557514 | 0.03060531 |
| 535 | 0.041085208 | 0.030560256 |
| 536 | 0.040797064 | 0.03006389 |
| 537 | 0.041257322 | 0.030668237 |
| 538 | 0.040610296 | 0.030040682 |
| 539 | 0.041278057 | 0.029169312 |
| 540 | 0.040911226 | 0.029855726 |
| 541 | 0.040595945 | 0.02918373 |
| 542 | 0.042377485 | 0.029198135 |
| 543 | 0.039844855 | 0.028926177 |
| 544 | 0.040222405 | 0.029041564 |
| 545 | 0.040557664 | 0.02896974 |
| 546 | 0.040948557 | 0.029918984 |
| 547 | 0.041229136 | 0.02972593 |
| 548 | 0.042018793 | 0.029700507 |
| 549 | 0.040926872 | 0.029586768 |
| 550 | 0.040638456 | 0.030291714 |
| 551 | 0.041457873 | 0.030528575 |
| 552 | 0.042017 | 0.030977213 |
| 553 | 0.041651907 | 0.03110746 |
| 554 | 0.041411748 | 0.031308639 |
| 555 | 0.043001153 | 0.031844072 |
| 556 | 0.041683419 | 0.031503805 |
| 557 | 0.041847941 | 0.031712862 |
| 558 | 0.042180993 | 0.031085786 |
| 559 | 0.042966746 | 0.031697395 |
| 560 | 0.041901694 | 0.032766419 |
| 561 | 0.041052921 | 0.032209208 |
| 562 | 0.042547549 | 0.032859811 |
| 563 | 0.043019746 | 0.031171994 |
| 564 | 0.042869082 | 0.031870075 |
| 565 | 0.04259555 | 0.031539041 |
| 566 | 0.042495922 | 0.031772912 |
| 567 | 0.041849738 | 0.03096196 |
| 568 | 0.042757629 | 0.031304522 |
| 569 | 0.042385588 | 0.031173599 |
| 570 | 0.040329248 | 0.029757283 |
| 571 | 0.041741617 | 0.029138568 |
| 572 | 0.041959 | 0.02928782 |
| 573 | 0.040394795 | 0.028463193 |
| 574 | 0.042377246 | 0.028834891 |
| 575 | 0.042868462 | 0.029326084 |
| 576 | 0.041217724 | 0.028643115 |
| 577 | 0.040870023 | 0.028252116 |
| 578 | 0.04152183 | 0.028335837 |
| 579 | 0.042072267 | 0.028919726 |
| 580 | 0.041743821 | 0.029298395 |
| 581 | 0.041806895 | 0.029681861 |
| 582 | 0.040685735 | 0.030320858 |
| 583 | 0.043016536 | 0.030965203 |
| 584 | 0.042402937 | 0.031996206 |
| 585 | 0.04392674 | 0.032806366 |
| 586 | 0.044501271 | 0.035278456 |
| 587 | 0.04229412 | 0.035422192 |
| 588 | 0.044236652 | 0.037092348 |
| 589 | 0.04453953 | 0.039432637 |
| 590 | 0.045403689 | 0.042120811 |
| 591 | 0.045720725 | 0.043050508 |
| 592 | 0.046721386 | 0.044569175 |
| 593 | 0.04658936 | 0.046035872 |
| 594 | 0.046856197 | 0.046439476 |
| 595 | 0.046639376 | 0.050515218 |
| 596 | 0.047224996 | 0.050214893 |
| 597 | 0.047826425 | 0.051104707 |
| 598 | 0.047684723 | 0.055223016 |
| 599 | 0.048337422 | 0.058213776 |
| 600 | 0.048084498 | 0.05737416 |
| 601 | 0.048056722 | 0.055091687 |
| 602 | 0.04834153 | 0.058816123 |
| 603 | 0.047162448 | 0.060133037 |
| 604 | 0.048829402 | 0.059604177 |
| 605 | 0.048451209 | 0.06028036 |
| 606 | 0.049314163 | 0.062367752 |
| 607 | 0.04864508 | 0.064056712 |
| 608 | 0.047748293 | 0.064284482 |
| 609 | 0.049373015 | 0.062747999 |
| 610 | 0.049654837 | 0.062923108 |
| 611 | 0.04873937 | 0.059863284 |
| 612 | 0.048195016 | 0.067000662 |
| 613 | 0.049368953 | 0.06636147 |
| 614 | 0.049471526 | 0.066356177 |
| 615 | 0.0509815 | 0.065962632 |
| 616 | 0.048364103 | 0.067198216 |
| 617 | 0.049258066 | 0.068526301 |
| 618 | 0.047925207 | 0.066248105 |
| 619 | 0.047651389 | 0.064842825 |
| 620 | 0.049918572 | 0.069186304 |
| 621 | 0.048079999 | 0.06930868 |
| 622 | 0.049001919 | 0.067431619 |
| 623 | 0.047876525 | 0.068410691 |
| 624 | 0.048913258 | 0.066037747 |
| 625 | 0.049283261 | 0.069188854 |
| 626 | 0.049833981 | 0.06884971 |
| 627 | 0.048520845 | 0.068593605 |
| 628 | 0.050626431 | 0.072924701 |
| 629 | 0.049001529 | 0.069368645 |
| 630 | 0.049823194 | 0.065875529 |
| 631 | 0.049922336 | 0.070392798 |
| 632 | 0.049238223 | 0.064130262 |
| 633 | 0.049091246 | 0.068703876 |
| 634 | 0.049004327 | 0.069478775 |
| 635 | 0.050113703 | 0.069644791 |
| 636 | 0.04935265 | 0.071714894 |
| 637 | 0.048524722 | 0.065578319 |
| 638 | 0.049033574 | 0.068908878 |
| 639 | 0.049706978 | 0.07082453 |
| 640 | 0.048870373 | 0.067637064 |
| 641 | 0.049378341 | 0.070513241 |
| 642 | 0.047531488 | 0.065338289 |
| 643 | 0.049938511 | 0.067545633 |
| 644 | 0.048420131 | 0.070092771 |
| 645 | 0.048394814 | 0.066225652 |
| 646 | 0.049045058 | 0.067036677 |
| 647 | 0.0471389 | 0.063334021 |
| 648 | 0.048276563 | 0.064587084 |
| 649 | 0.049566595 | 0.067005786 |
| 650 | 0.048757214 | 0.066615875 |
| 651 | 0.048086512 | 0.064861617 |
| 652 | 0.049770716 | 0.071548828 |
| 653 | 0.048497445 | 0.065184075 |
| 654 | 0.051019438 | 0.064323894 |
| 655 | 0.047818361 | 0.069356187 |
| 656 | 0.047395308 | 0.06791304 |
| 657 | 0.048211057 | 0.064465262 |
| 658 | 0.04925706 | 0.068522353 |
| 659 | 0.049243152 | 0.067570369 |
| 660 | 0.049306566 | 0.068635881 |
| 661 | 0.048296998 | 0.064357191 |
| 662 | 0.04946561 | 0.069889086 |
| 663 | 0.048651964 | 0.066769534 |
| 664 | 0.049580237 | 0.061861251 |
| 665 | 0.047312599 | 0.061431122 |
| 666 | 0.048177763 | 0.066106882 |
| 667 | 0.049260615 | 0.06726634 |
| 668 | 0.050070504 | 0.065101662 |
| 669 | 0.049296071 | 0.060405176 |
| 670 | 0.049355936 | 0.066502011 |
| 671 | 0.048166386 | 0.06328206 |
| 672 | 0.048552887 | 0.063599076 |
| 673 | 0.047215508 | 0.0630013 |
| 674 | 0.050010272 | 0.065692928 |
| 675 | 0.048251383 | 0.059550299 |
| 676 | 0.049073229 | 0.064339078 |
| 677 | 0.049158252 | 0.066160593 |
| 678 | 0.048586706 | 0.063461555 |
| 679 | 0.049408432 | 0.066435927 |
| 680 | 0.048615398 | 0.063448449 |
| 681 | 0.048136575 | 0.06465852 |
| 682 | 0.049240503 | 0.065996535 |
| 683 | 0.048443426 | 0.068781814 |
| 684 | 0.049338138 | 0.064378941 |
| 685 | 0.048442599 | 0.067303664 |
| 686 | 0.047661626 | 0.060606162 |
| 687 | 0.048758079 | 0.06567617 |
| 688 | 0.048292359 | 0.062784951 |
| 689 | 0.048590671 | 0.064714098 |
| 690 | 0.049590564 | 0.062694093 |
| 691 | 0.047962726 | 0.063263871 |
| 692 | 0.050139198 | 0.061484326 |
| 693 | 0.048513909 | 0.063766816 |
| 694 | 0.048005757 | 0.062790064 |
| 695 | 0.048824793 | 0.06817339 |
| 696 | 0.048301867 | 0.062689082 |
| 697 | 0.048157362 | 0.065482695 |
| 698 | 0.047640732 | 0.059392198 |
| 699 | 0.048021823 | 0.062982676 |
| 700 | 0.049106955 | 0.063966326 |

**Supplementary Video 1 (Related to Figure 1**). One representative simulation of paired excitation and emission photons traveling. Left: Initial ballistic photon flow of a 488 nm excitation beam incident from the optical fiber. Right: Diffuse propagation of the elicited 561 nm emissions captured by the same fiber.

**Figure S1.**
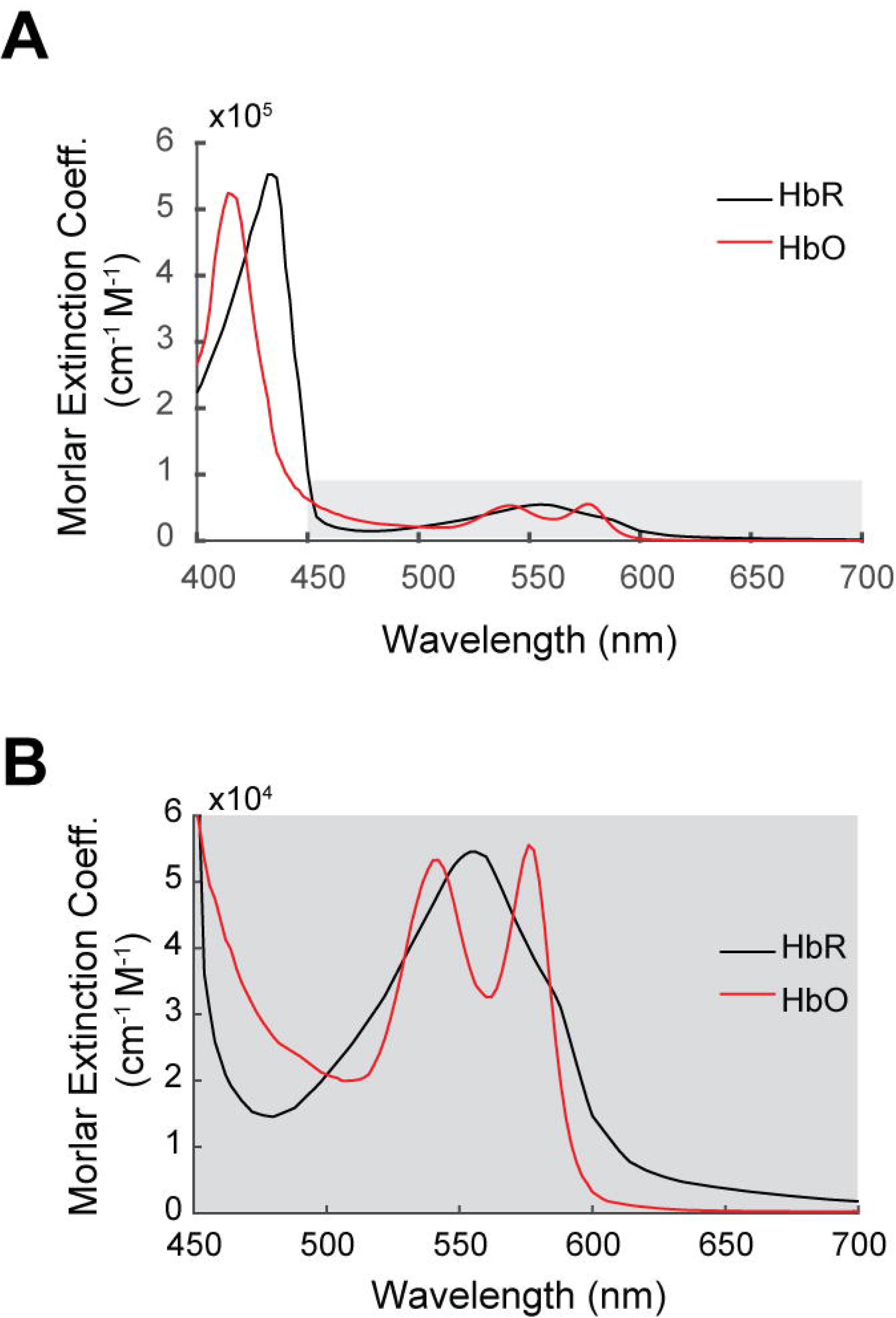
(**Related to Figure 1)**. Molar extinction coefficients of HbO and HbR as a function of wavelength. **(A)** Broad-band presentation. **(B**) The range commonly used to measure fluorescent sensor activity.

**Figure S2.**
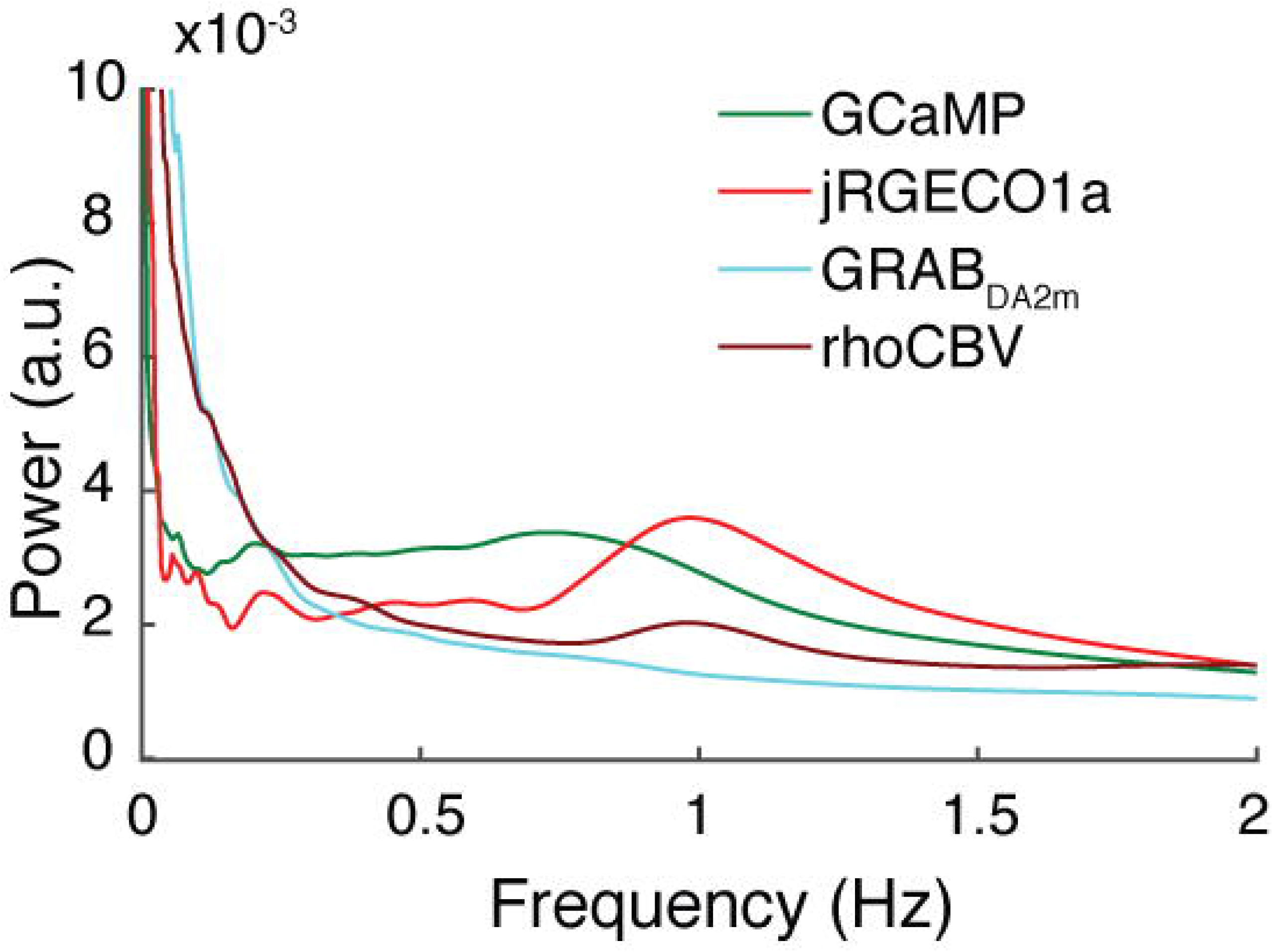
(Related to Figure 1). Power spectral distribution of several commonly used fluorescent sensors. CBV were measured by intravenously injecting a long-circulating red fluorescent dye, Rhodamine B (rhoCBV). Data were acquired continuously to generate time-courses and then converted to power spectra density using Fourier transform.

**Figure S3.**
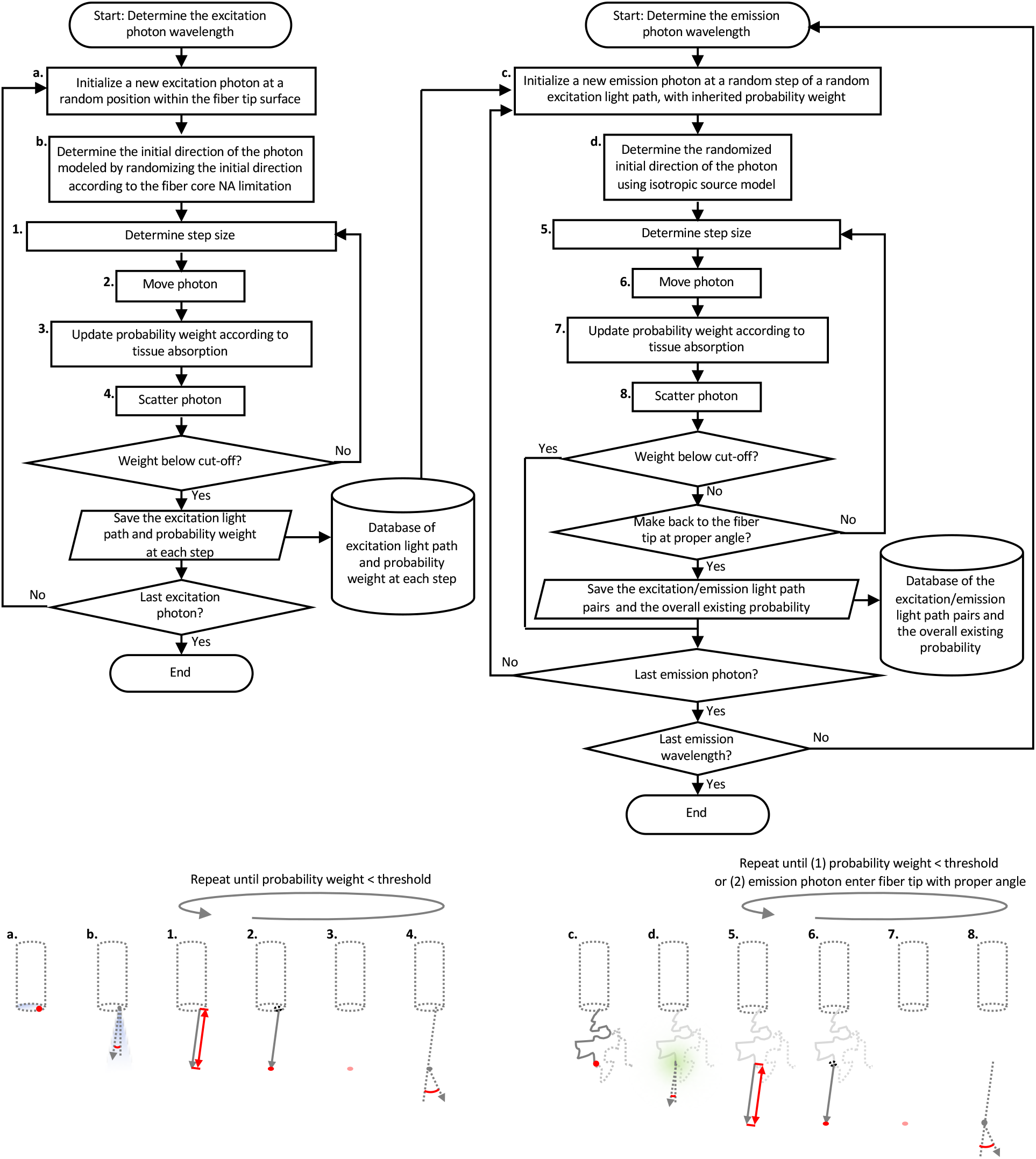
(**Related to Figure 1**). Light path simulation flow chart.

**Figure S4.**
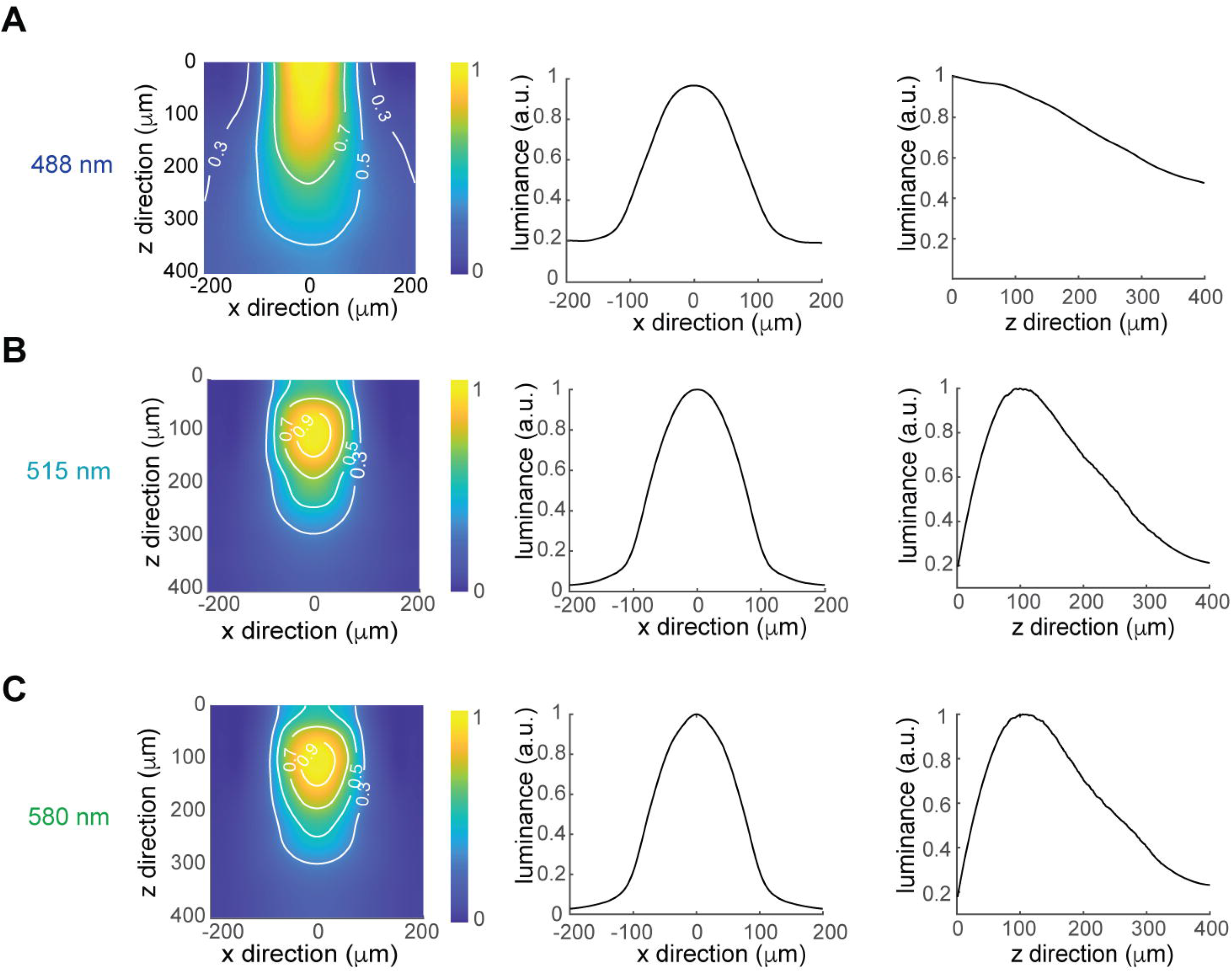
(Related to Figure 1). Monte Carlo simulation results. **(A)** Illuminating distribution of the excitation 488 nm light. Left showing the spatial probability map, and middle and right showing normalized luminance-distance profiles in x and z directions. **(B)** Receptive probability of the 515 nm emission light. Left showing the spatial probability map, and middle and right showing normalized luminance-distance profiles in x and z directions. **(C)** Receptive probability of the 580 nm emission light. Left showing the spatial probability map, and middle and right showing normalized luminance-distance profiles in x and z directions. Color bar indicates probability from 0 to 1.

**Figure S5.**
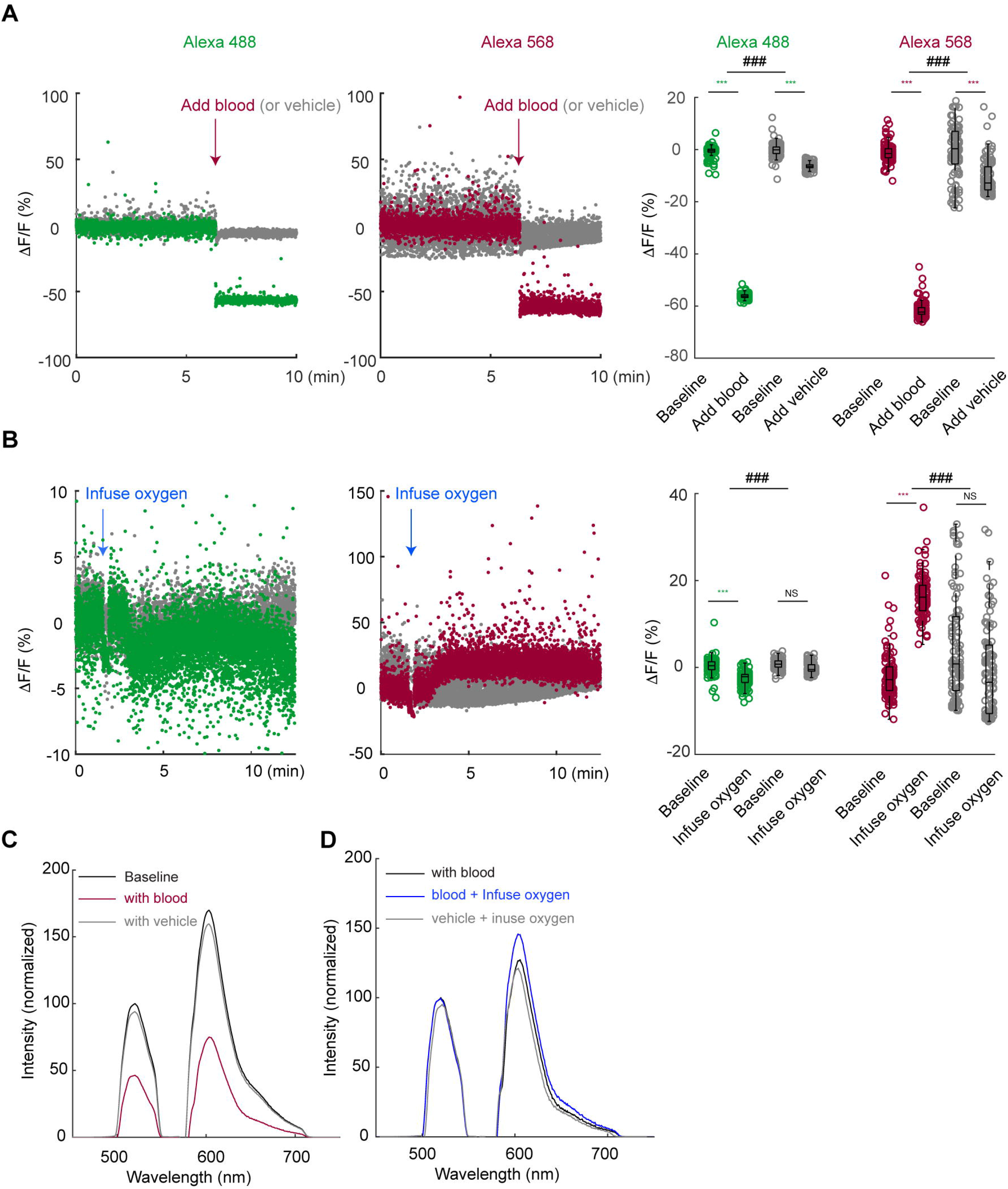
Ex vivo quantification of Hb-absorption effects on photometry recordings. (**A**) Δ*F/F* changes before and after adding 1 mL of fresh mouse venous blood or 1 mL vehicle (0.9% saline) into 10 mL saline solution containing 10 μL Alexa 488 and 10 μL Alexa 568 fluorescent dyes. Adding saline decreased the concentrations of Alexa 488 and 568, which caused the slight but significant reduction in signal intensity. Error bars represent standard deviation. *** *p* < 0.001, *t*-test (n = 100 time points). ### *p* < 0.001, two-way ANOVA (n = 100). (**B**) Δ*F/F* changes before and after 100% oxygen infusion into the solution containing blood or vehicle and fluorescent dye mixture. Error bars represent standard deviation. *** *p* < 0.001, *t*-test (n = 100 time points). ### *p* < 0.001, two-way ANOVA (n = 100). (**C**) Normalized intensity spectra of the fluorescent solution mixture before and after adding blood or vehicle. (**D**) Normalized intensity spectra of the blood-fluorescent solution mixture, or vehicle-fluorescent solution mixture, before and after oxygen infusion.

**Figure S6.**
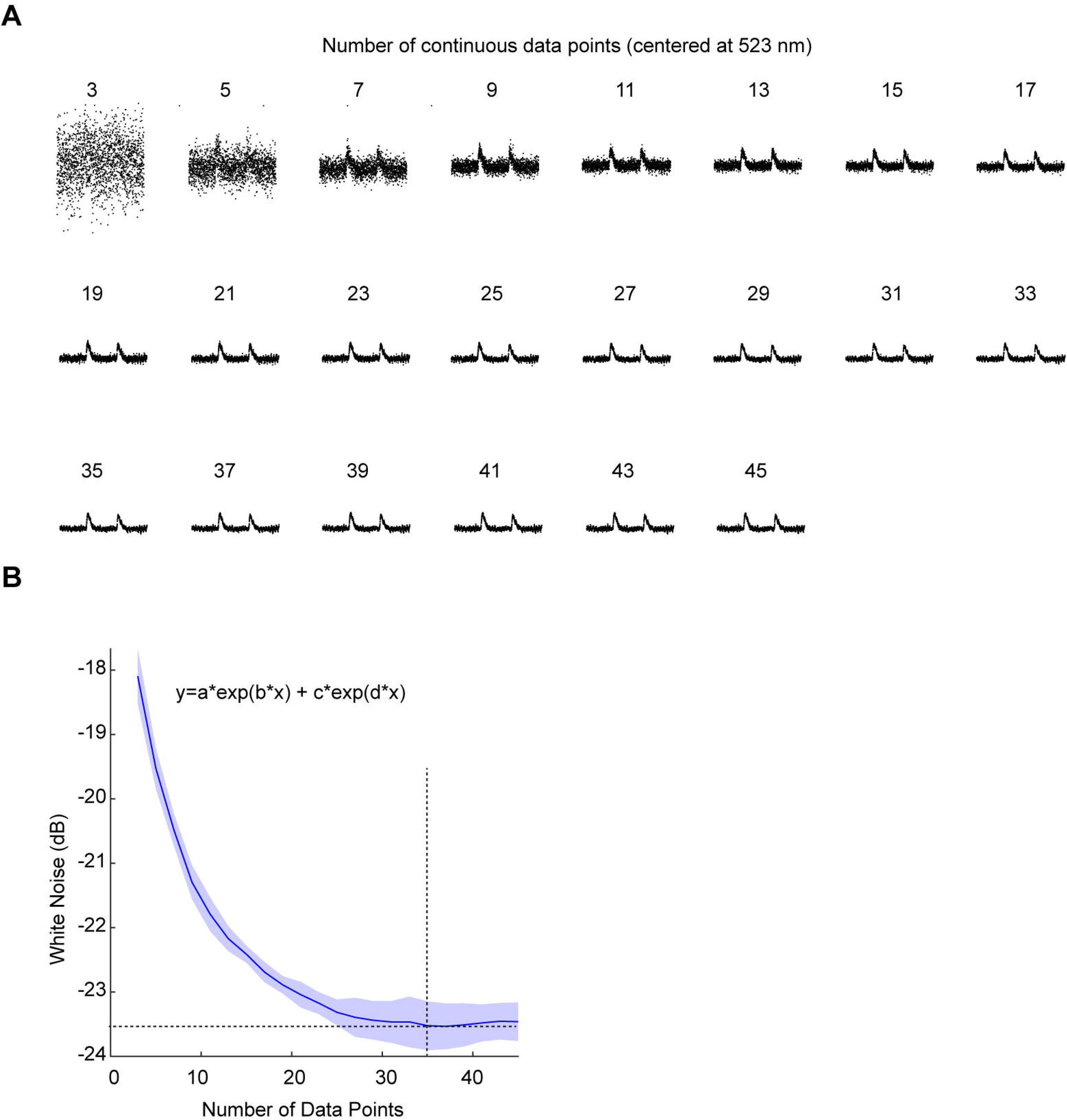
(Related to Figure 2). Number of continuous spectral datapoints needed to derive reliable hemoglobin concentration changes. (**A**) Representative Δ*C_HbO_* time-courses derived from activity-independent EYFP data using different number of spectral datapoints ranging from 3 to 45 (centered at 523 nm). (**B**) Reduction of white noise was observed with increased number of datapoints. The derived Δ*C_HbO_* exhibited minimized white noise when the number of datapoints reaching 35.

**Figure S7.**
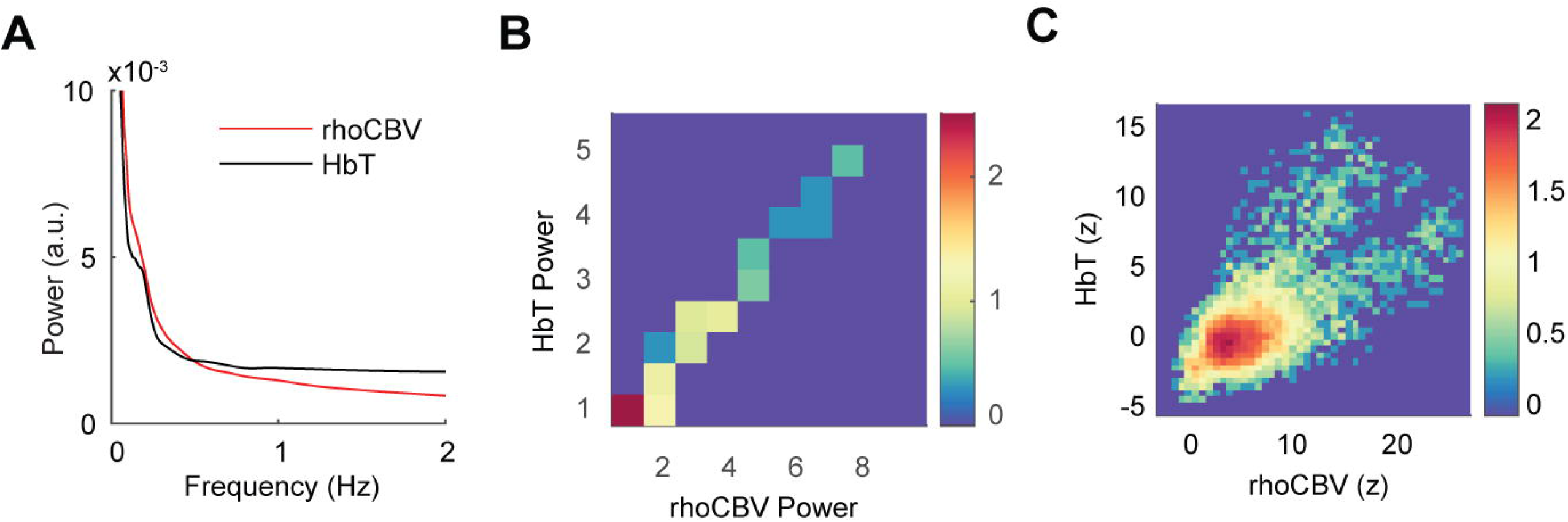
(Related to Figure 2). Comparison between corrected rhoCBV and HbT derived from EYFP signals. (**A**) Power spectral distribution. (**B**) Two-dimensional histogram summarizing rhoCBV and HbT power. Each datapoint in the frequency series is assigned to a two-dimensional matrix to make the histogram. Correlation is significant at the 0.001 level (Pearson correlation, *r* = 0.98). The color bar indicates counts on the log scale. (**C**) Two-dimensional histogram summarizing z-scores of rhoCBV and HbT. Each datapoint in the timecourses is assigned to a two-dimensional matrix to make the histogram. Correlation is significant at the 0.001 level (Pearson correlation, *r* = 0.66). The color bar indicates counts on the log scale.

**Figure S8.**
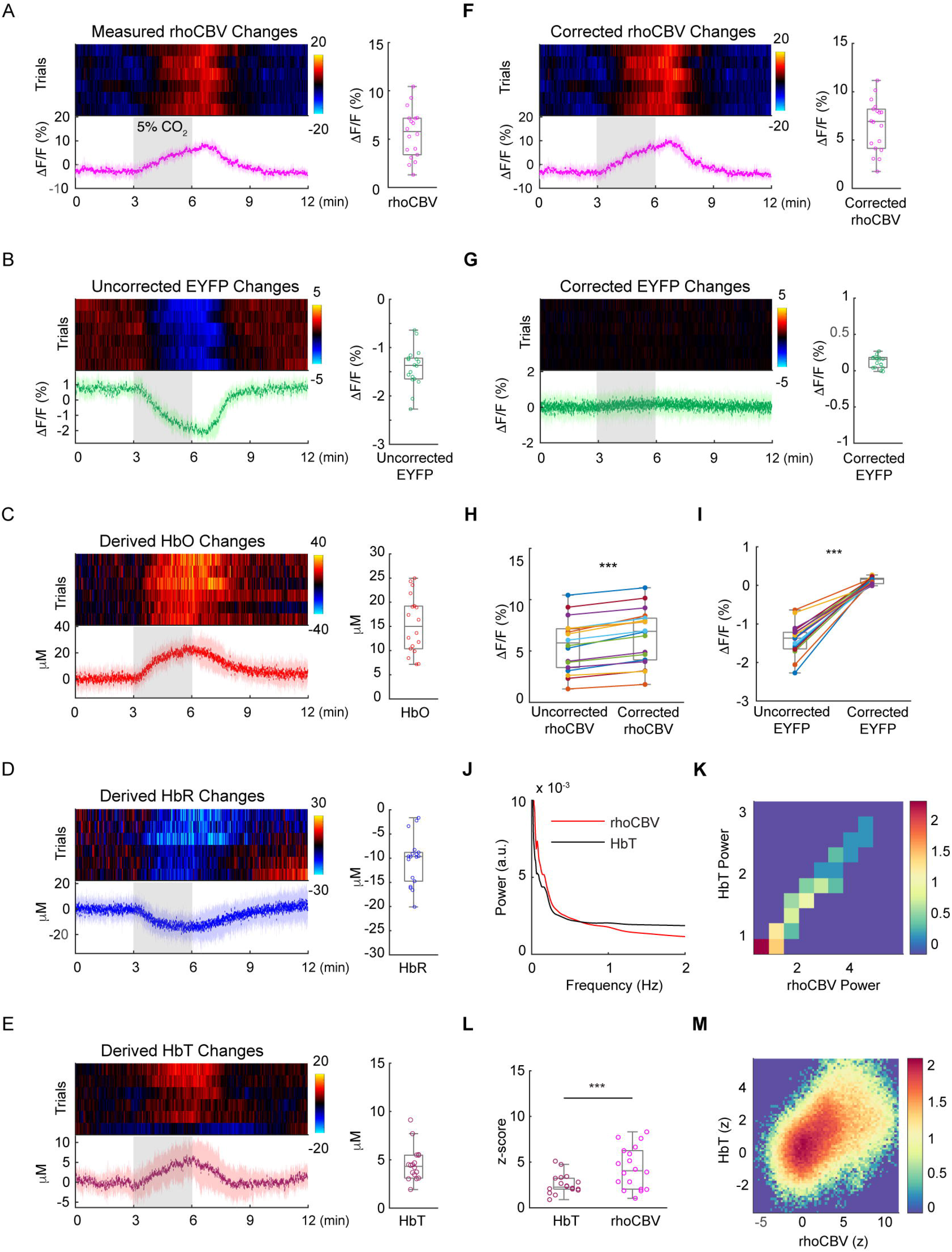
(Related to Figure 2). Fiber-photometry in the S1 with EYFP during hypercapnia (5% CO_2_). (**A-G**) Peri-event heatmaps showing repeated trials and average response time-courses of the changes in measured rhoCBV, measured EYFP, derived HbO, derived HbR, derived HbT, corrected rhoCBV, and corrected EYFP, respectively. The gray-shaded segment indicates infusion of 5% CO_2_. All color bars use the same unit as the time-course Y axis. Error bars represent standard deviation. (**H, I**) Correction of the contaminated rhoCBV and EYFP signals, respectively, across all subjects and all trials (n=5, total trials =18; *** indicates *p* < 0.001, paired *t*-test). (**J**) Power spectral distribution. (**K**) Two-dimensional histogram summarizing rhoCBV and HbT power. Each datapoint in the frequency series is assigned to a two-dimensional matrix to make the histogram. Correlation is significant at the 0.001 level (Pearson correlation, *r* = 0.98). The color bars indicates counts on the log scale. (**L**) rhoCBV exhibited higher sensitivity than derived HbT. Error bars represent standard deviation. *** indicates p < 0.001 (paired *t*-test). (**M**) Two-dimensional histogram summarizing rhoCBV and HbT z-scores. Each datapoint in the timecourses is assigned to a two-dimensional matrix to make the histogram. Correlation is significant at the 0.001 level (Pearson correlation, *r* = 0.50). The color bars indicates counts on the log scale.

**Figure S9.**
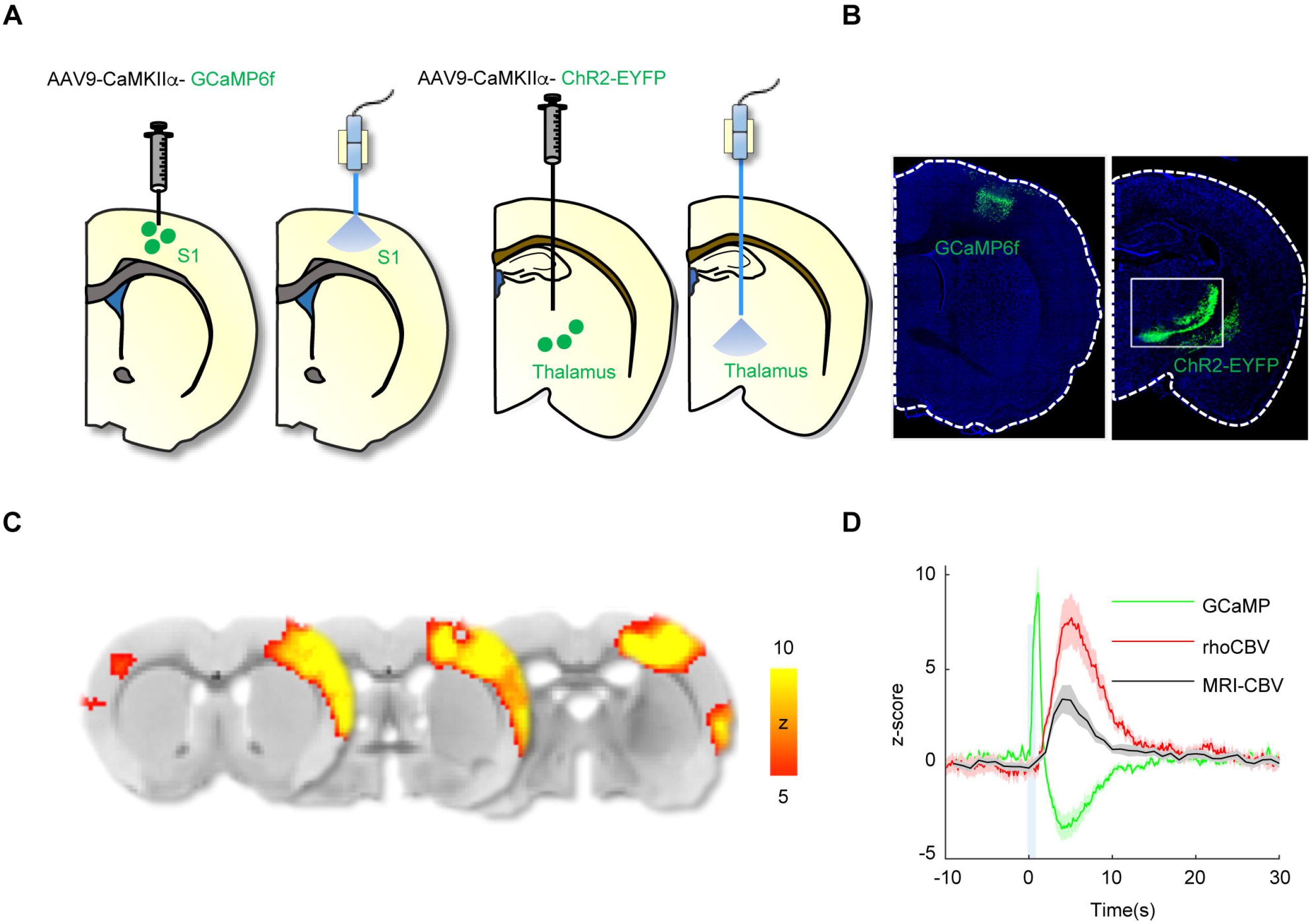
(Related to Figure 3). Simultaneous fMRI and fiber-photometry recording showing optogenetic-evoked CBV-fMRI, GCaMP and rhoCBV changes in the S1. (**A**) GCaMP and channelrhodopsin-2 (ChR2) were virally expressed using AAV under the CaMKIIα promoter in the S1 and ventroposterior (VP) thalamus, respectively. Rhodamine B was intravenously injected to achieve CBV measurement. (**B**) Confocal images showing GCaMP expression. (**C**) MRI-CBV activation maps in response to optogenetic stimulation of VP. (**D**) Uncorrected GCaMP, rhoCBV, and MRI-CBV time-courses (n=3, total trials =33 trials). After a short optogenetic stimulation of VP (60Hz, 10mW power, 0.5 ms pulse width for 1 s), uncorrected GCaMP signal exhibited a strong undershoot with signal pattern mirrored the rhoCBV and MRI-CBV data, indicative of significant Hb absorption *in vivo*.

**Figure S10.**
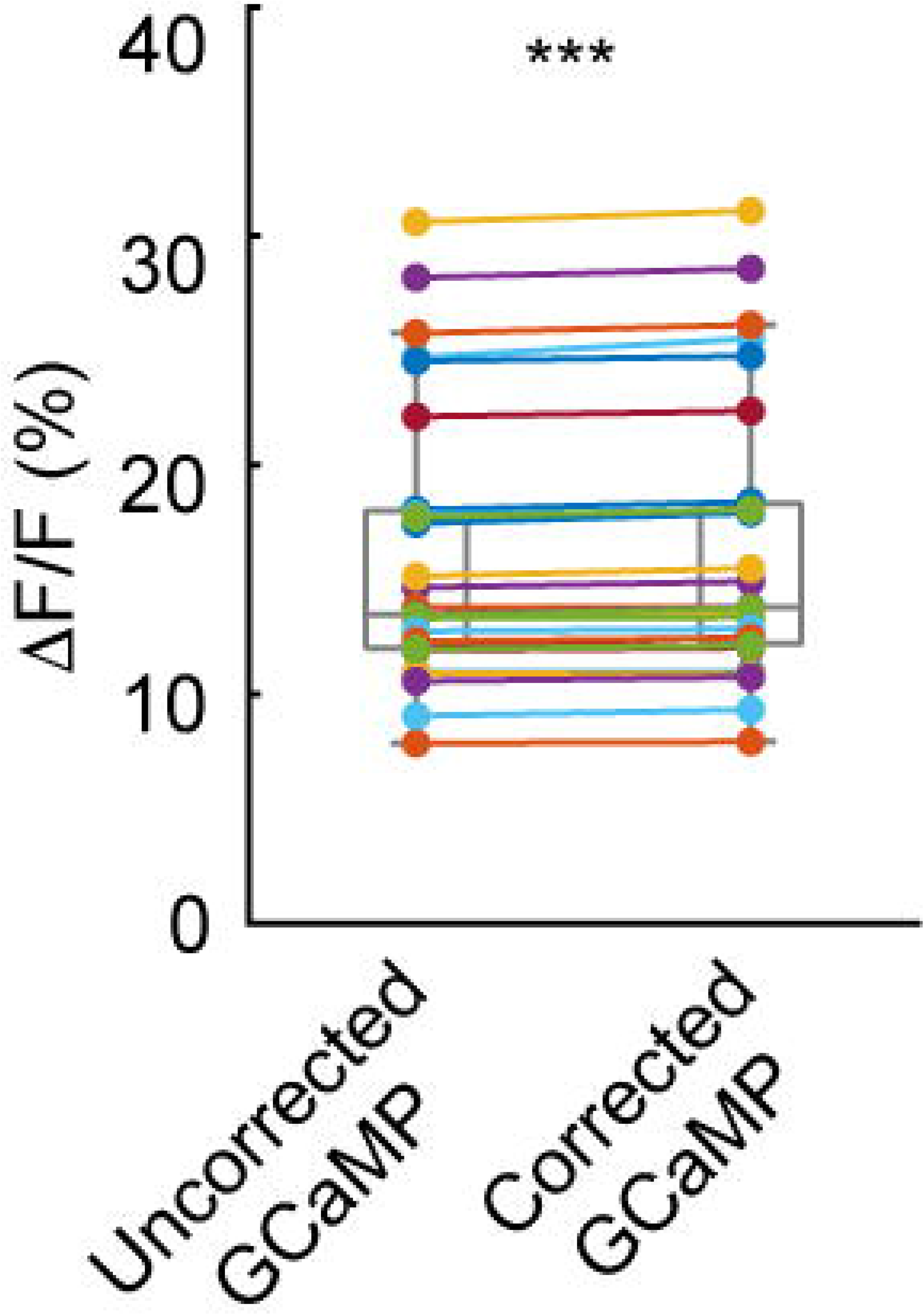
(Related to Figure 3). Comparison of the peak GCaMP responses before and after Hb-absorption correction. The peak amplitude of GCaMP responses is significantly higher in the corrected GCaMP compared with uncorrected GCaMP. (*** indicates *p*<0.001, paired *t*-test, n=27)

**Figure S11.**
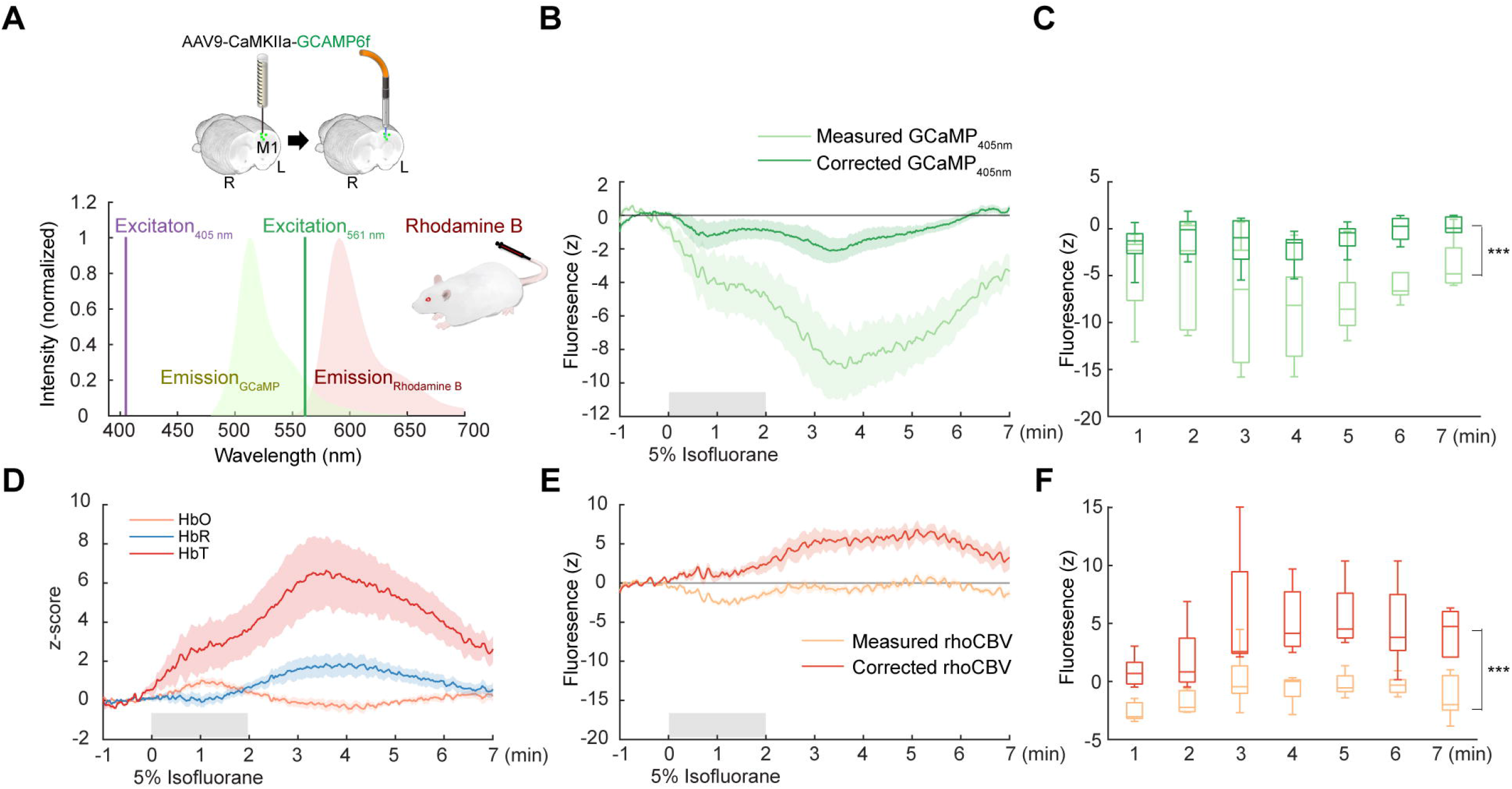
(Related to Figure 4). Hb-absorption effects on GCaMP signal measurement during 5% isoflurane challenges. **(A)** Preparation for fiber-photometry measurement of GCaMP and Rhodamine B signals in the left M1. Ca^2+^-independent GCaMP signal (GCaMP_405_) was excited by a 405 nm laser and used for hemodynamic correction. rhoCBV signals were measured as described in **Figure 2**. **(B)** Measured and corrected GCaMP_405_ time-courses before, during, and after 2-min 5% isoflurane challenges (baseline = 0.5% isoflurane). The gray-shaded segment indicates the isoflurane period. Error bars represent standard deviation. **(C)** Scatter plots of measured and corrected GCaMP_405_ signals at 1, 2, 3, 4, 5, 6, and 7 min after the onset of the isoflurane challenge (n = 5, *p* < 0.001, two-way ANOVA). **(D)** Derived HbO, HbR, and HbT time-courses before, during, and after the isoflurane challenge. **(E)** Measured and corrected rhoCBV time-courses before and after the isoflurane challenge. Read-out from the Rhodamine B could be misinterpreted without hemodynamic correction. **(F)** Scatter plots of measured and corrected rhoCBV signals at 1, 2, 3, 4, 5, 6, and 7 min after the onset of the isoflurane challenge (n = 5, *p* < 0.001, two-way ANOVA).

**Figure S12.**
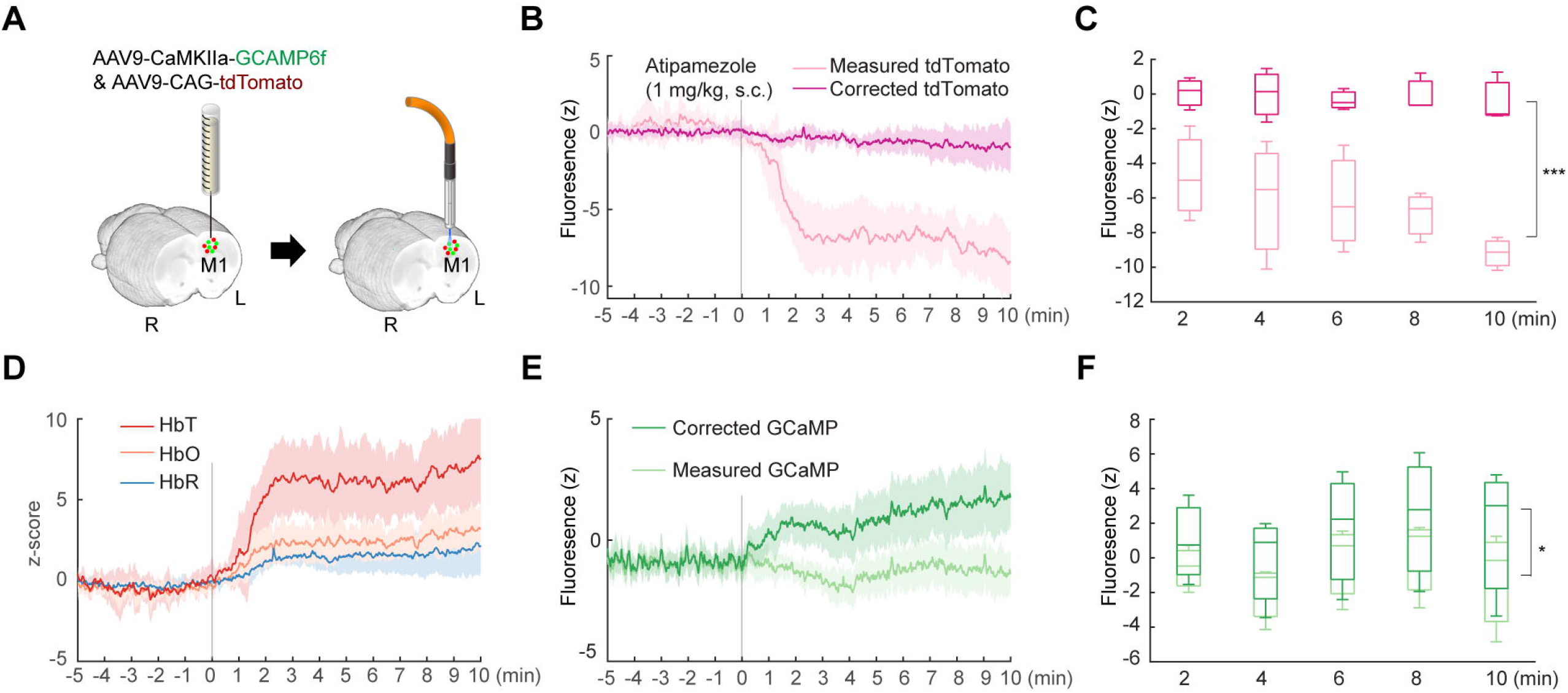
(Related to Figure 4). Hb-absorption effects on GCaMP signal measurement during pharmacological studies. (A) Fiber-photometry measurement of GCaMP and tdTomato signals in the left M1. (B) Measured and corrected tdTomato time-courses before and after administering an α_2_ adrenergic antagonist Atipamezole (1 mg/kg, s.c., Zoetis, Parsipanny, NJ). (C) Scatter plots of measured and corrected tdTomato signals at 2, 4, 6, 8, and 10 min after the atipamezole administration (n = 3, *p* < 0.001, two-way ANOVA). (D) Derived HbO, HbR, and HbT time-courses before and after the atipamezole administration. Increased CBV might result from the redistribution of blood supply to active brain regions (Bekar et al., 2012). (E) Measured and corrected GCaMP time-courses before and after the atipamezole administration. the increased GCaMP activity in the cortex would have been otherwise unchanged without hemodynamic correction. (F) Scatter plots of measured and corrected GCaMP signals at 2, 4, 6, 8, and 10 min after the atipamezole administration (n = 3, *p* < 0.05, two-way ANOVA).

**Figure S13.**
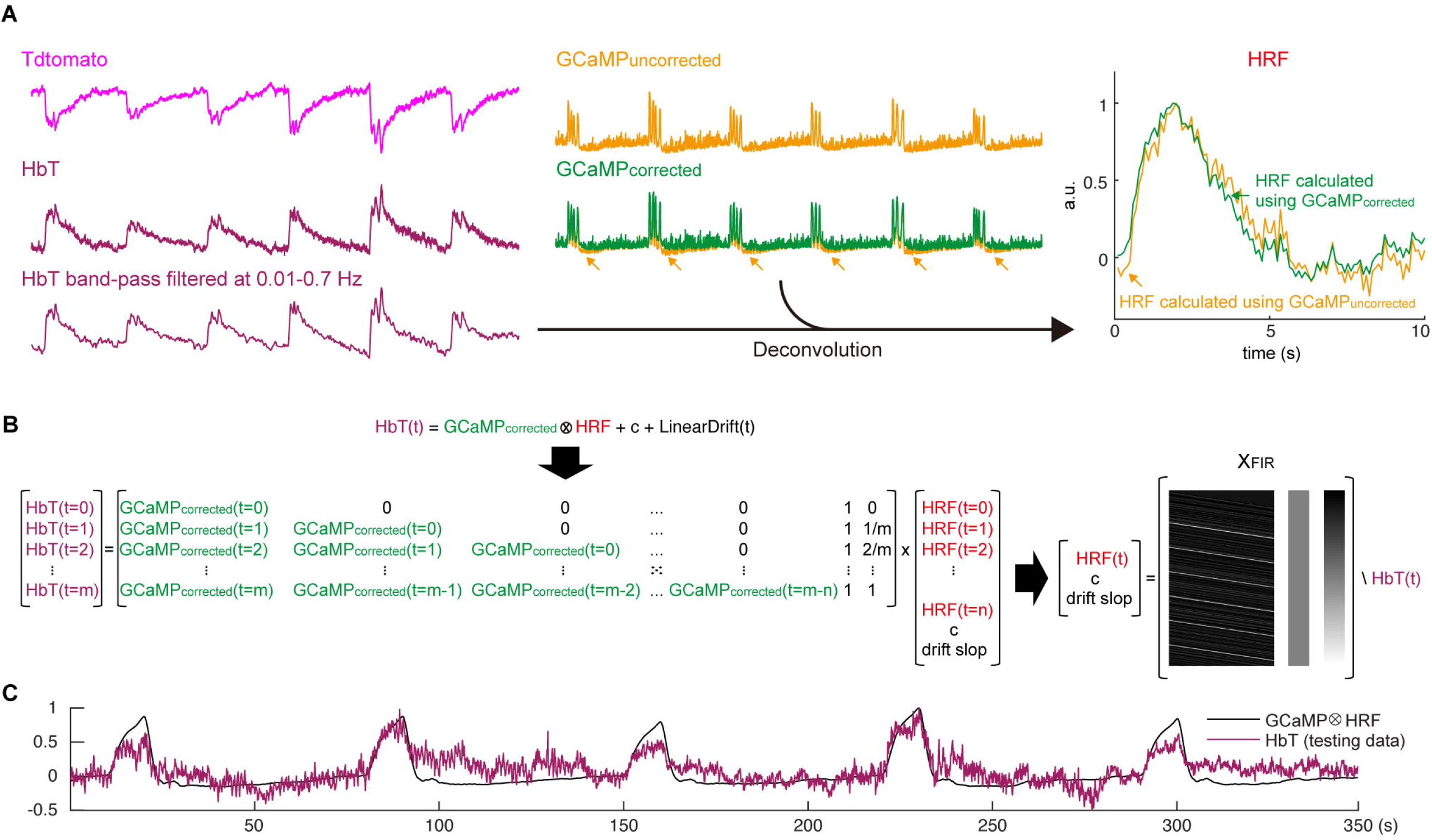
(Related to Figure 5). HRF calculation and validation. (**A**) Pipeline for estimating HRF from tdTomato-derived HbT and GCaMP signals. Yellow arrows indicate post-stimulus undershoot caused by Hb absorption. Note that using uncorrected GCaMP caused an HRF amplitude shift at t=0. (**B**) Strategy to deconvolve HRF form preprocessed HbT and GCaMP signals. (**C**) Validation of the HRF using a separate dataset naïve to HRF calculation. HRF was convolved with GCaMP data to predict HbT changes (black trace). High correlation was observed between the predicted and tdTomato-derived HbT (purple trace).

**Figure S14.**
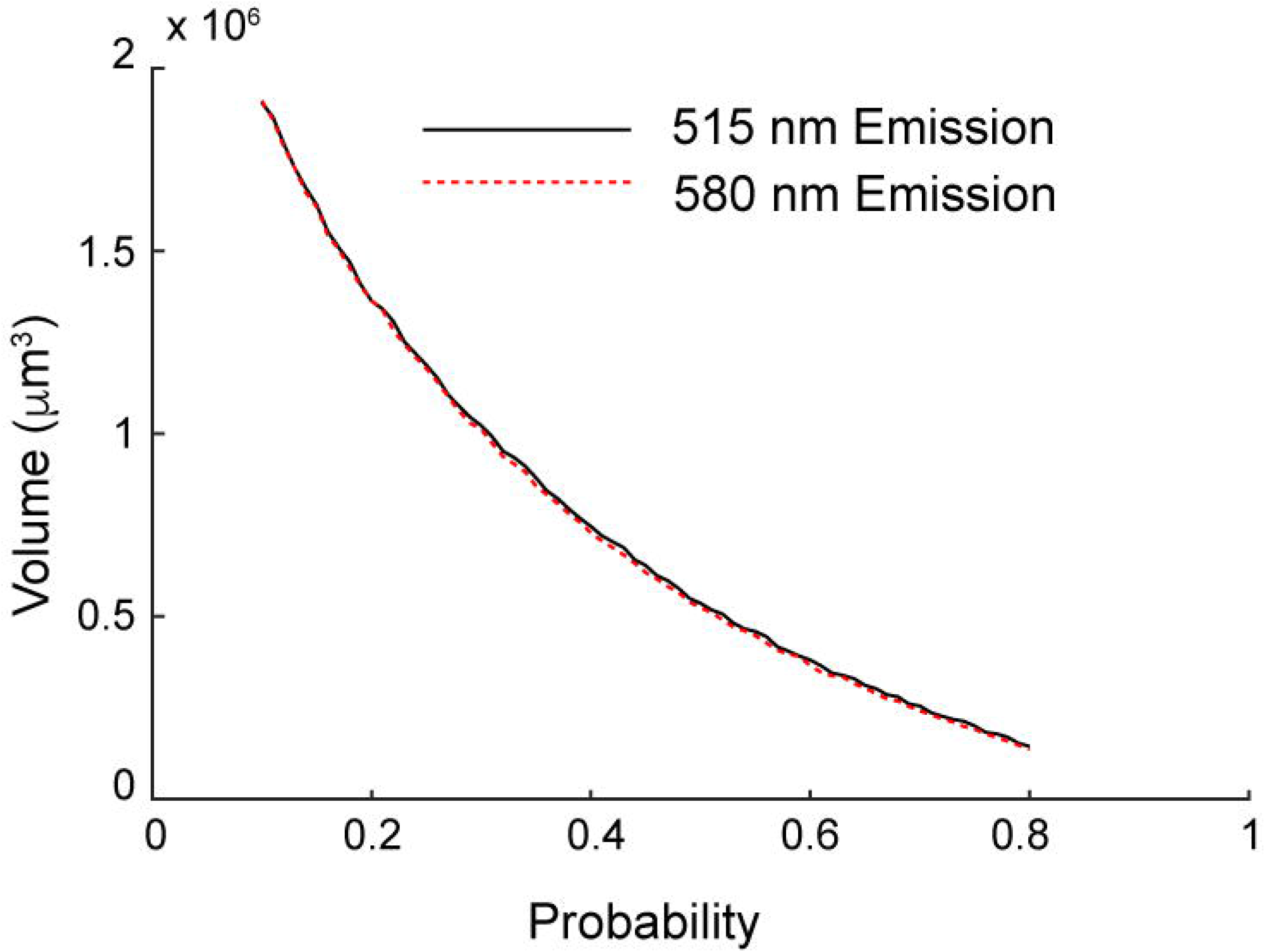
Simulated detection volume and its projected probability.

**Figure S15.**
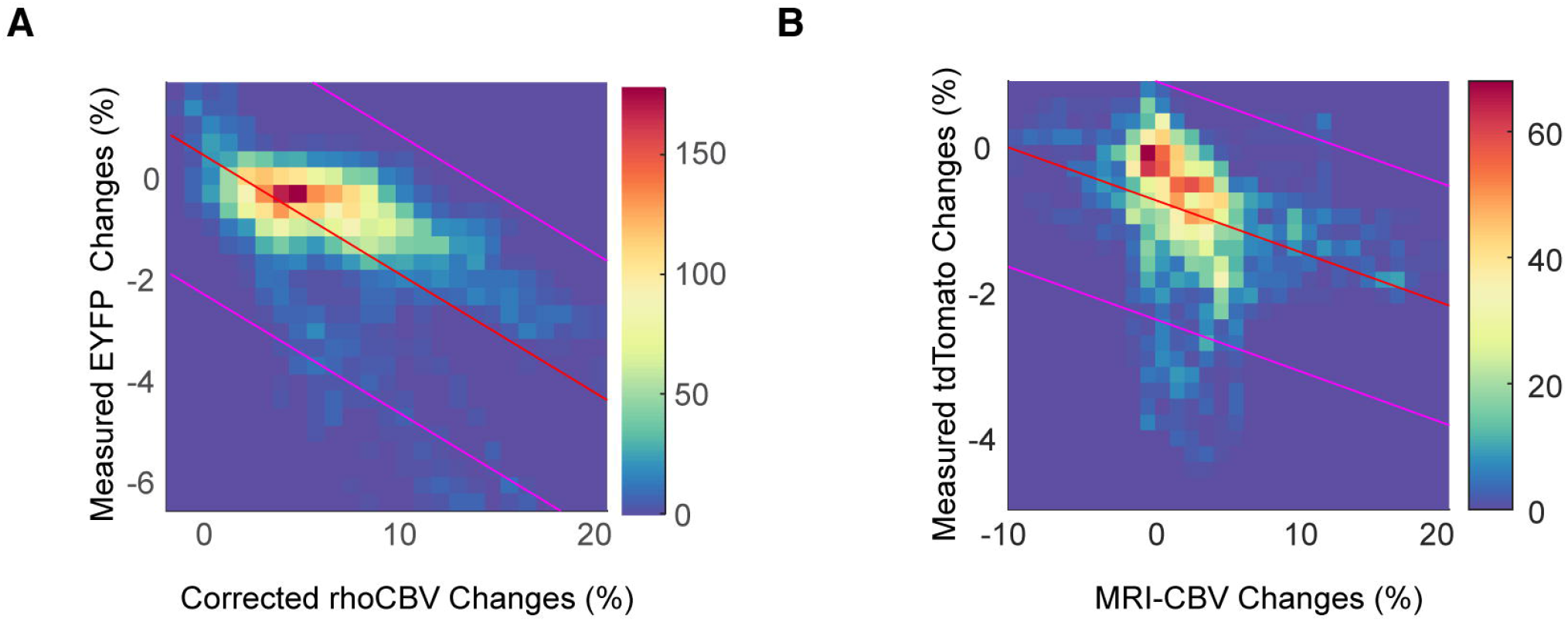
Signal drop in measured EYFP and tdTomato signals were correlated to the CBV changes. (**A**) Two-dimensional histogram summarizing % changes of corrected rhoCBV and measured EYFP signal. Each datapoint in the time-courses is assigned into a two-dimensional matrix to make the histogram. Correlation is significant at the 0.001 level (Pearson correlation, *r* = −0.62). Color bars indicate datapoint counts. Linear fit (red line) and 95% confidence interval (pink lines) are also shown. (**B**) Two-dimensional histogram summarizing % changes of MRI-CBV and measured tdTomato signal. Each datapoint in the time-courses is assigned into a two-dimensional matrix to make the histogram. Correlation is significant at the 0.001 level (Pearson correlation, *r* = −0.33). Color bars indicate datapoint counts. Linear fit (red line) and 95% confidence interval (pink lines) are also shown.

**Figure S16.**
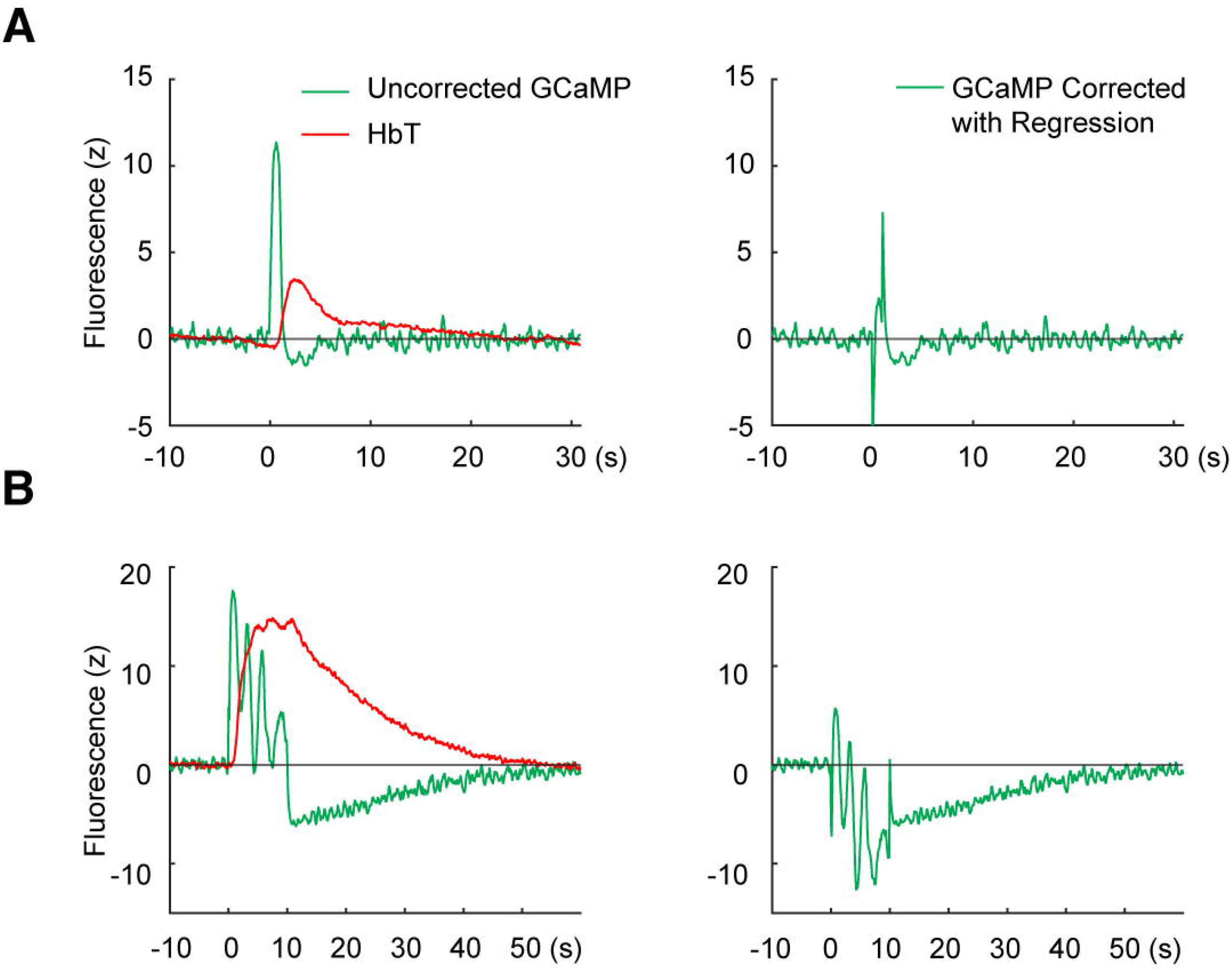
Correcting GCaMP signals by regressing out the HbT. **(A)** short (1 s) stimulation **(B)** long (10 s) stimulation.

**Figure S17.**
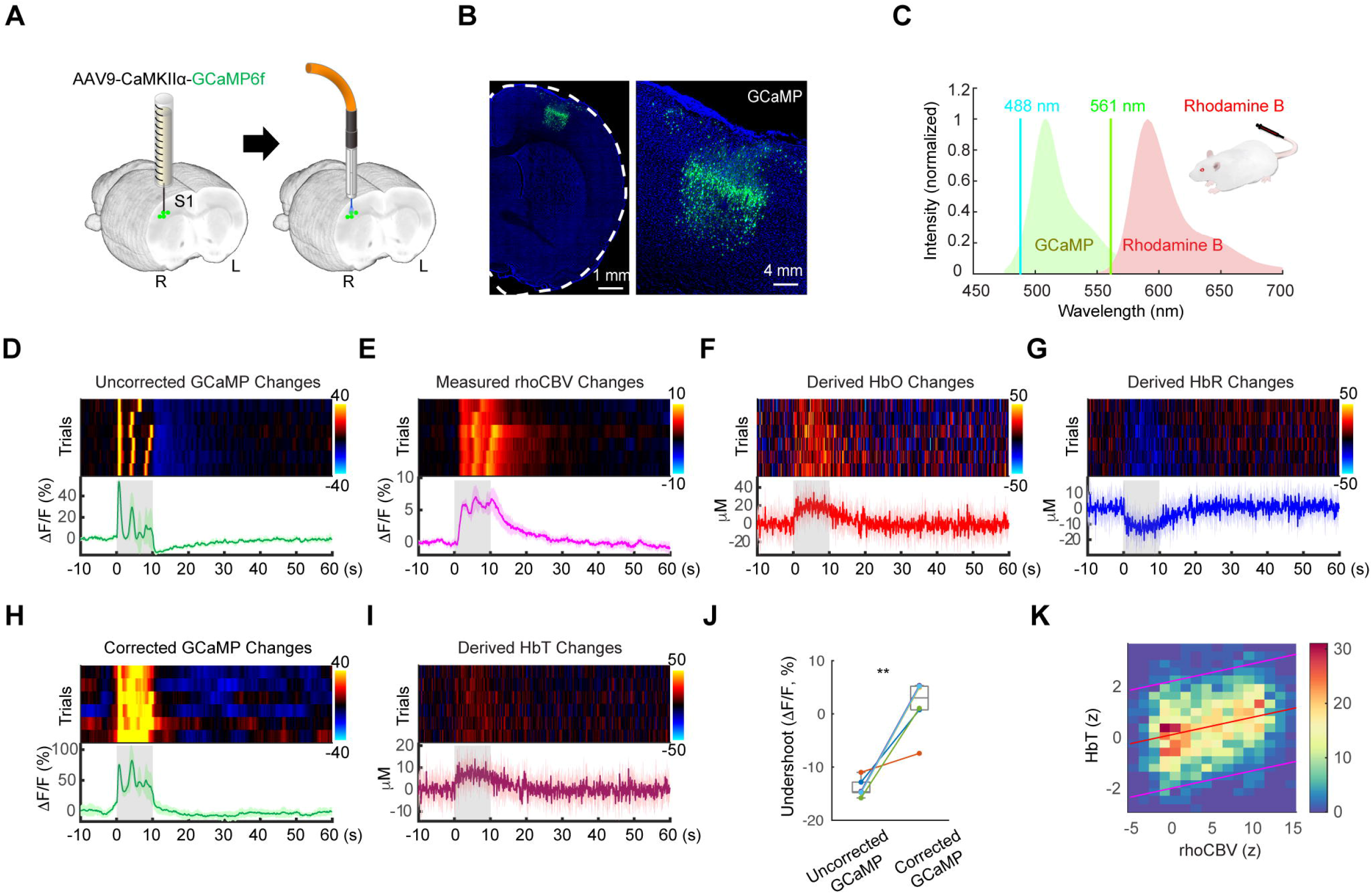
Quantification of Hb-absorption effects using residue signals of fiber-photometry. (**A**) Preparation for fiber-photometry measurement of GCaMP signal in the S1 of the forelimb region. (**B**) Confocal images showing GCaMP expression. (**C**) Simultaneous dual-spectral photometry recording of GCaMP and rhoCBV. (**D-I**) Peri-stimulus heatmaps showing repeated trials and average response time-courses of the changes in uncorrected GCaMP, measured rhoCBV, derived HbO, derived HbR, corrected GCaMP, and derived HbT, respectively. The gray-shaded segment indicates forepaw-stimulation period. All color bars use the same unit as the time-course Y axis. Error bars represent standard deviation. (**J**) Correction of the post-stimulus dip of GCaMP signals (n = 2, total trials = 6; ** indicates *p* < 0.01, paired *t*-test). (**K**) Two-dimensional histogram summarizing rhoCBV and HbT z-scores. Each datapoint in the timecourses is assigned into a two-dimensional matrix to make the histogram. Correlation is significant at the 0.001 level (Pearson correlation, *r* = 0.28). Colorbar indicates the number of datapoints. Linear fit (red line) and 95% confidence interval (pink lines) are also shown.

